# Chimpanzees use numerous flexible vocal sequences with more than two vocal units: A step towards language?

**DOI:** 10.1101/2021.02.03.429517

**Authors:** Cédric Girard-Buttoz, Emiliano Zaccarella, Tatiana Bortolato, Angela D. Friederici, Roman M. Wittig, Catherine Crockford

**Author notes:** These authors contributed equally (Cédric Girard-Buttoz, Emiliano Zaccarella).

## Abstract

A major question in evolutionary science is how did language evolve? Syntax, as the core of language, combines meaning-bearing units (words) into hierarchical structures, thereby creating new meanings. Some other mammals and birds combine meaning-bearing vocalisations, but no documented examples exist of non-human animals combining more than two meaning-bearing vocalisations. Was the two-unit threshold only surpassed in the hominid lineage? Here, we examine the positional patterning of vocal sequences of chimpanzees. We analysed 4826 vocal utterances of 46 wild adult female and male chimpanzees. We found a flexible system with 390 multi-unit vocal sequences, some showing positional or transitional regularities. Two-unit pairs embedded in three-unit sequences predictably occurred either in head or tail positions, and co-occurred with specific other elements. The capacity to organise vocal output beyond the two-unit level may thus exist in species other than humans and could be viewed as an important evolutionary step towards language.

## Introduction

A major conundrum in evolutionary science has been reconstructing the evolution of language^1–5^. Given that language does not fossilize, a key line of research has been comparative, comparing the communication systems of other animals with that of humans. Some would argue this approach has only increased the enigma, as animal communication, especially in non-human primates (hereafter primates), is considered to show very limited communication capacities^6,7^. Primates are thought to be highly constrained in their vocal production by a fixed vocal repertoire with very limited learning potential^8,9^. This is in stark contrast with human syntax. Only humans have the sophisticated system of syntax, where meaning-bearing units (words) can be endlessly combined in hierarchical structures through recursion that create infinite new meanings^10,11^; but how and when did this capacity evolve?

Comparative studies have examined the capacities of other animals to both comprehend and produce vocal sequences. A sequence is broadly defined as the production of two or more different types of vocal units within a short time from each other^12^. In terms of comprehending or processing sequences, auditory discrimination tasks using familiarization-discrimination paradigms on artificially generated rule-based sequences have however shown that some primates (cotton-top tamarins and macaques) appear to be sensitive to local sequence violations generated by grammars of regular kinds, i.e., [(AB)^2^]^13,14^. This suggests that primates might possess some organizational principles applying over pattern regularities between neighbouring units. Note that these tasks encode patterning but do not encode meaning as sounds are selected so as to be neutral and devoid of context. Crucially, such positional principles appear to go beyond simple adjacency rules to include the capacity to detect sequence violations across non-adjacent relationships between distinct sound^15,16^ or visual^17^ units.

In terms of producing sequences, comparative studies examining production in non-vocal domains, such as learning signing or use of symbol charts, show that non-human primates fail to approach human abilities to generate complex sequences beyond the two- “word” or “unit” level. Studies on natural production of gestures highlight the presence of sequences in great ape gestural communication^18–20^. Yet, whether these sequences follow some ordering rules and convey semantic information remains to be established^19^. This supports the discontinuity claim that primates might lack combinatorial grammar systems when conveying meaningful information^21–23^.

In terms of examining natural vocal repertoires, playback experiments, where single calls or call sequences are played through a loud speaker whilst controlling for certain social or environmental criteria, demonstrate that some calls convey specific information to conspecifics, such as a type of predator^24^ or the need to recruit others to join^25^. These are called *meaning-bearing vocal units*. Some animals combine two of such meaning-bearing units together resulting in modifications to meaning of at least one of the single units (e.g., forest monkeys^26^; Japanese tits^27^; and pied babblers^25^).

In terms of natural vocal production, recent and insightful reviews have tried to relate meaning-bearing call combinations to linguistic principles^26,28–30^. Miyagawa & Clarke^29^ have, however, suggested that, in primates, meaning-bearing units are not produced in sequences that are longer than two units. Reviews on call sequences production across birds and mammals also do not provide examples of meaning bearing units being emitted in longer than two-unit sequences^26,28^. It should be noted that the production of vocal sequences with more than two different vocal units are common across animals but are rarely demonstrated to contain individual meaning-bearing units^12,28^. Such longer vocal sequences generally occur in species’ song (e.g., songbirds^31^, humpback whales^32^, and monogamous primates^33^) or in their long distance calls (e.g., new world monkeys^34,35^). Here, changes in sequence diversity or ordering generally function to emphasize individual quality or identity in courtship and resource defence contexts^12^ but units in the sequence are not independently meaning-bearing.

In sum, on the one hand, sequence *processing* studies suggest that primates might own principles of combination which are sensitive to positional information, independent of meaning, and which might be used to establish sequential relationships across multiple sound units. On the other hand, spontaneous *vocal production* studies suggest a limit to the length of call sequences that contain meaning-bearing units at the “two-unit level”. This apparent discrepancy in sequence processing versus production capacities may arise because selection for flexible processing has been greater than for flexible production, suggesting an ancestral dissociation between the two^36^. Alternatively, the apparent discrepancy may be due to limited assessment of longer vocal sequences and related methodological difficulties^12^. In primates, for example, call sequences comprising more than two types of meaning-bearing units have been reported^34,37^, but a minority of call sequences have been analysed quantitatively for either pattern variation or contextual specificity (but see ^38^).

A promising approach to test for the potential to combine more than two meaning-bearing units in non-human animals is to determine the potential for a pairing system. Here pairing refers to the frequent adjacent production of two independent meaningful vocal units that are used in a sequence with additional vocal units. Additional units can occur either before – e.g., δ + [*νω*] or ε + [*νω*] – or after the pair – e.g., [*νω*] + δ or [*νω*] + ε, creating longer vocal sequences. A pairing system can extend to the joining of different pairs within the same sequence: [*νω*] + [δε] (see ^12^).

To test for the potential for such a “pairing” system, we require a non-human species with natural vocal output that fits at least three criteria: 1) the species emits vocal sequences that contain more than two different vocal units, 2) the units included in such sequences have the potential to be meaning-bearing; and 3) the vocal sequences are longer than two-units and comprise single vocal units which are ordered predictably and consistently within the sequence A first step in identifying the latter is to establish which sequences are produced above chance level (i.e., more frequently than by random juxtaposition of vocal units). A second step is to establish if vocal units produced singularly are also regularly and predictably emitted in these two-unit non-random sequences. A third step is to determine whether such two-unit sequences are in turn predictably reused within several different three-unit sequences.

One species that fulfils the first two requirements is the chimpanzee. Chimpanzees use numerous different vocal sequences not only in their loud calls (pant-hoot^39^) but across most parts of their vocal repertoire^40^. Vocal sequences make up towards 50% of their call output and regularly involve sequences of more than two different types of vocal units^40,41^. Chimpanzee single vocal units have a relatively high degree of context-specificity across a relatively broad range of contexts: alarm, recruit, hunt, food, submissive greeting, rest, travel, nest^42^. Whilst combinations of two meaning-bearing vocal units are not uncommon across mammals and birds, these tend to be limited to alarm or recruitment contexts^24,28^.

The goal of our study is to investigate whether the vocalizations produced by chimpanzees exhibit the features of a potential pairing system. More specifically, we aim to establish whether sequences extending beyond two-units reuse pairs of calls used in two-vocal unit sequences and whether the ordering of vocal units within sequences follows a certain regularity. To reach this goal we first extracted and reported the use of single units as well as all the vocal sequences with two or more (up to ten) vocal units, using a broad full-vocal repertoire analysis. The specific criteria to define the time frame between the two or more units produced within a sequence varies across studies^12^. In our study, we used a conservative 1s maximum gap between calls.

In a second step, we specifically concentrated on two- and three-vocal unit sequences, which represent about 80% of the vocal sequences recorded in this study. We first determined the potential positional bias for single and two-unit pairs to occur in head or tail positions within two-unit and three-unit sequences, respectively, using a Bayesian binomial approach. Second, we assessed which specific two- and three-vocal units occur above chance level within the sample using randomizations. This step was necessary to select sequences that were regularly produced, and to focus our analysis on the ordering of vocal units within these sequences specifically. Accordingly, as a last step, we examined transitional relationships between units within these above chance sequences, to determine the likelihood for single or paired units to follow or to be followed by other specific units. These three analyses were conducted to assess whether the sequences produced by chimpanzees in our study are just a random juxtaposition of single vocal units, or if they rather show some ordering rules/ non-random order among the individual vocal units.

We use 900.8 h of data from 46 wild mature chimpanzees from Taï National Park, Ivory Coast, fully habituated to human observers, from three communities. We used a systemic whole repertoire approach where vocalisations were continuously recorded during focal animal sampling.

We used a call classification procedure, training listeners to classify calls according to sound and spectrographic information, resulting in high inter-ratter reliability scores between coders. This method was proven to be the most accurate at classifying calls, especially in a noisy forest environment, as compared to semi-automatic classification of manually-extracted metrics or fully automatic call classification algorithms^12^. We also assessed the accuracy of the classifications using acoustic analysis on a subset of calls and discriminant function analysis. To limit classification problems arising from the highly acoustically graded chimpanzee repertoire, we chose to classify broad call categories that show consensus across studies and chimpanzee populations (reviewed in ^42^; **Table 1**, **Fig. S1**) and the acoustic properties of which discriminate well in discriminant function analysis (Grawunder et al. In revision). Single calls, such as grunts and hoos, are frequently combined with a voiced inhalation (pant), producing a string of repetitions of alternations of pant + another vocalization (**Figure S1b and S2**). The use of unpanted or panted forms can result in contextual shifts. Single grunts, for example, are predominantly emitted at food, whereas panted grunts are predominantly emitted as a submissive greeting vocalization^39,42^. It could be argued that panted forms of vocalisations are in themselves a call sequence. However, as they can be context-specific, are produced independently, are common in the chimpanzee repertoire, and can themselves be combined with other vocalisations, we have defined these as separate vocal units.

**Table 1.** Discrimination of chimpanzee call types.

| Call type | Abbreviation | Fundamental Frequency (f0) | Shape of f0 | Noisiness | Single/panted | Occurrence in the repertoire |
| --- | --- | --- | --- | --- | --- | --- |
| Grunt | GR | 70-700 Hz | Variable | Noisy | Single | 2087 (27.1%) |
| Panted grunt | PG | 100-200 Hk | Variable | Noisy | Panted | 853 (11.1 %) |
| Hoo | HO | 200-700 Hz | Flat | Highly tonal | Single | 1561 (20.3%) |
| Panted hoo | PH | - | Flat | Highly tonal | Panted | 1015 (13.2%) |
| Bark | BK | 600-2000 Hz | Dome | Variable | Single | 337 (4.4%) |
| Panted bark | PB | 600-2000 Hk | Dome | Variable | Panted | 549 (7.1 %) |
| Pant | PN | 100-200 Hz (if visible) | Flat (if visible) | Noisy | Panted | 402 (5.2%) |
| Scream | SC | 800-2000 Hz | Flat mid section | Variable | Single | 312 (4.1%) |
| Panted scream | PS | 800-2000 Hz | Flat mid section | Variable | Panted | 317 (4.1%) |
| Non-voiced sounds (lip smack, raspberry) | NV | - | - | Noisy | Single | 184 (2.39%) |
| Whimper | WH | 350-1300 Hk | Flat | Highly tonal | Single | 78 (1.0%) |
| Panted roar | PR | 200-300 Hk, with bands below F0 (<100 Hz) | Flat | Noisy | Panted | 1 (0.01%) |

In the chimpanzee repertoire, variants of the same call, such as a hoo, are emitted in different contexts, such that the acoustic variants are context-specific, and elicit different behavioural responses during playback experiments from receivers^43^. However, it may be that hoos share a common function, such as to coordinate activities like resting and travelling^42,44^. Likewise, although screams are emitted in a range of contexts, and some can be discriminated acoustically from others^45^, they likely share an overarching function to recruit others^42,44^. Hence, for simplicity, in this study, we do not distinguish between different call variants but classify all hoos, as ‘hoo’, all screams as ‘scream’, and so on. Thus, we used a simplified classification that did not include call variants. This approach will, if anything, under-represent variation in vocal sequence use.

## Results

### Frequency distribution of utterances

Chimpanzees produce 12 different types of vocal units (i.e., different call types, see **Table 1** for the abbreviation of the name of these different call types as used in the result section). For this study, we recorded 4826 utterances from 46 mature chimpanzees (**Fig. 1**) i.e., older than 10 years of age. We define as mature individuals adult and subadult male and female chimpanzees. 3232 (67.2 %) of these utterances were single calls (2747 utterances as single vocal units and 485, c.a. 10%, in panted form). Single units did not comprise sequences of different vocal units (**Fig. 1**, **Table S1**). 1584 (32.8%) utterances were emitted as vocal sequences including 817 (16.9%) pairs (i.e., sequences with two vocal units) and 458 (9.5%), 170 (3.5%), 90 (1.9%), 29 (0.6%), 12 (0.2%), 4 (0.1%), 2 (0.04%) and 2 (0.04%) sequences comprising three, four, five, six, seven eight, nine and 10 vocal units respectively (**Fig. 1**, **Table S1**). The length of a sequence was determined based on the succession of two or more vocal units which were each different from the preceding unit. However, the same vocal unit could be repeated in the sequence when separated by at least one other vocal unit type (e.g., A_B_C_C was counted as a sequence of three vocal units, whereas A_C_B_C was counted as a sequence of four vocal units). From an average of 13.2 focal hours per individual, all individuals produced sequences with at least three vocal units. Two-thirds of the chimpanzees (32/46: 69.6%) produced sequences with five vocal units. Only females produced sequences with eight or more vocal units. Examples of sequences produced by chimpanzees are provided in **Fig. S3**.

**Figure 1:**
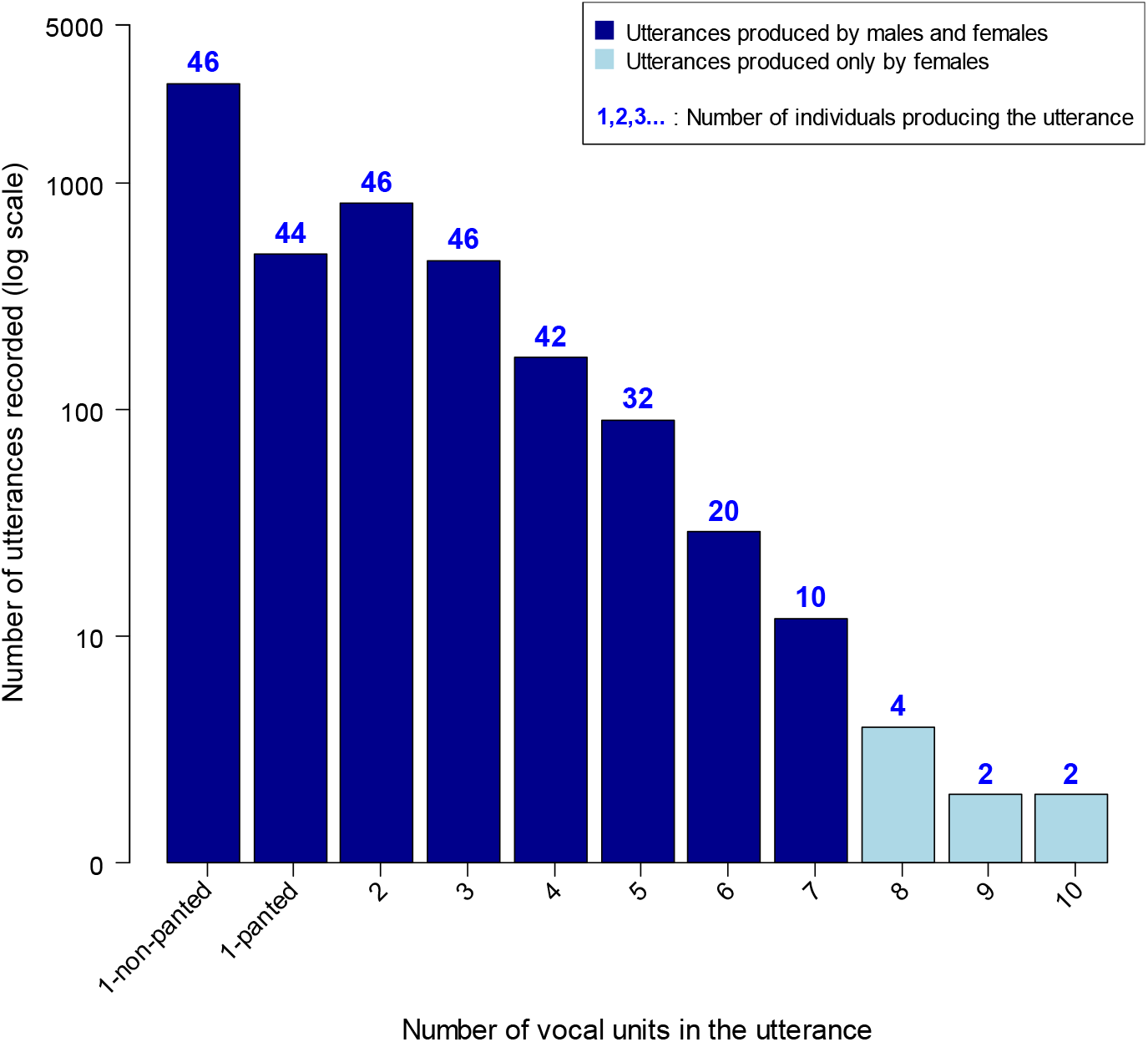
Frequency distribution of vocal sequences recorded with different lengths (i.e., different number of vocal units). The number of vocal units in the utterance is indicated on the x-axis. The y-axis depicts the number of recordings (log-transformed for ease of visualization). All utterances with one to seven vocal units (in blue) are produced by both sexes while utterances with eight or more vocal units (in light-blue) are only produced by females. The number on top of each bar in blue indicates the number of individuals producing utterances of this particular length.

### Vocal sequence analysis: two vocal units (pairs)

The chimpanzees produced 58 unique sequences with two vocal units (pairs, **Table S1**). The number of occurrences of each vocal unit in first and second positions within each of the two-unit sequences can be found in **Fig. 2A** (see also **Table S2**). We represent all the two-vocal unit sequences in the dataset as a combinatorial network, and show the corresponding combinatorial strength per unit (number of in/out connections) in **Fig. 2B-C** (see also **Fig. S4**). Within the network, PH appears to be the call that is used the most with all other calls in the dataset (Betweenness Centrality (BC) = 12.45). This suggests that PH might be a key call in the two-unit network and that it can be emitted flexibly in sequence with all other calls (HO=4.566, SC=4.116, GR=3.816, BK=3.5, PB=2.533, PG=1.115, WH=1.033, PS=0.666, PN=0.2).

**Figure 2:**
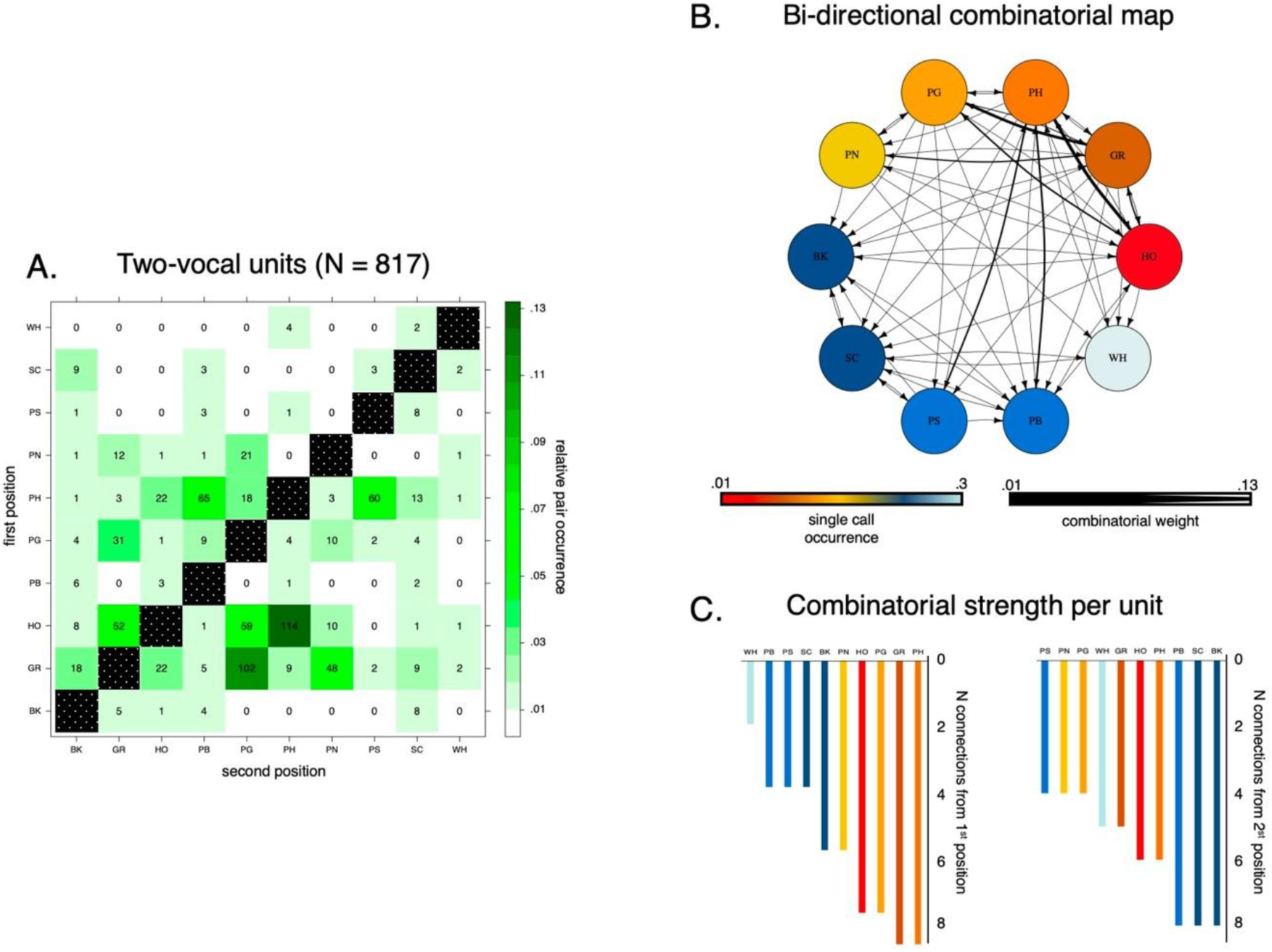
(A) Frequency distribution for two-unit vocal sequences with call units occurring in first position listed along the y-axis and call units occurring in second position listed along the x-axis. Colour gradients (white-to-dark-green) represent the relative occurrence of each pair within the two-vocal units’ set. In each cell, the absolute frequency count for each pair is conversely reported. (B) The two-vocal units’ combinatorial network with the ten units depicted as circled nodes and colour gradients (hot-to-cold) representing the number of times a certain unit is found in the two-vocal units’ set. This network was also used for the calculation of the Betweenness Centrality among the units. The size of the directional edges (arrows) expresses the number of times the specific two-vocal unit is found in the sample (thick-to-thin). (C) The number of different vocal units with which each call forms a pair in the sample as first unit (left) or second unit in the pair (right).

We first asked three questions concerning the organization of single units to form sequences of two vocal units (hereafter pairs). (1) Positional bias: Are the vocal units produced by chimpanzees biased towards the start or end position within a pair? (2) Specific ordering frequency: Do the pairs show fixed order frequencies that go beyond random juxtaposition of single vocal units? (3) Transitional bias: Do relationships between these units exist when forming sequences of two vocal units (i.e., call *ν* always follows call *ω*, or call *ν* always precedes call *ω*)?

#### Positional Bias

We used Bayesian binomial tests (see Methods and **Table 2A**) over the number of occurrences of each call in either first or second position within a pair. We found strong evidence that HO, GR and PH vocal units most reliably occured in the first position in two-unit sequences (**Fig. 3A** **and** **Table 2A**). Conversely, PG, PB, PS, SC, BK, and PN showed an opposite bias towards second position (**Table 2A**). We found no bias for WH, which however only occurred 13 times in total.

**Table 2.**
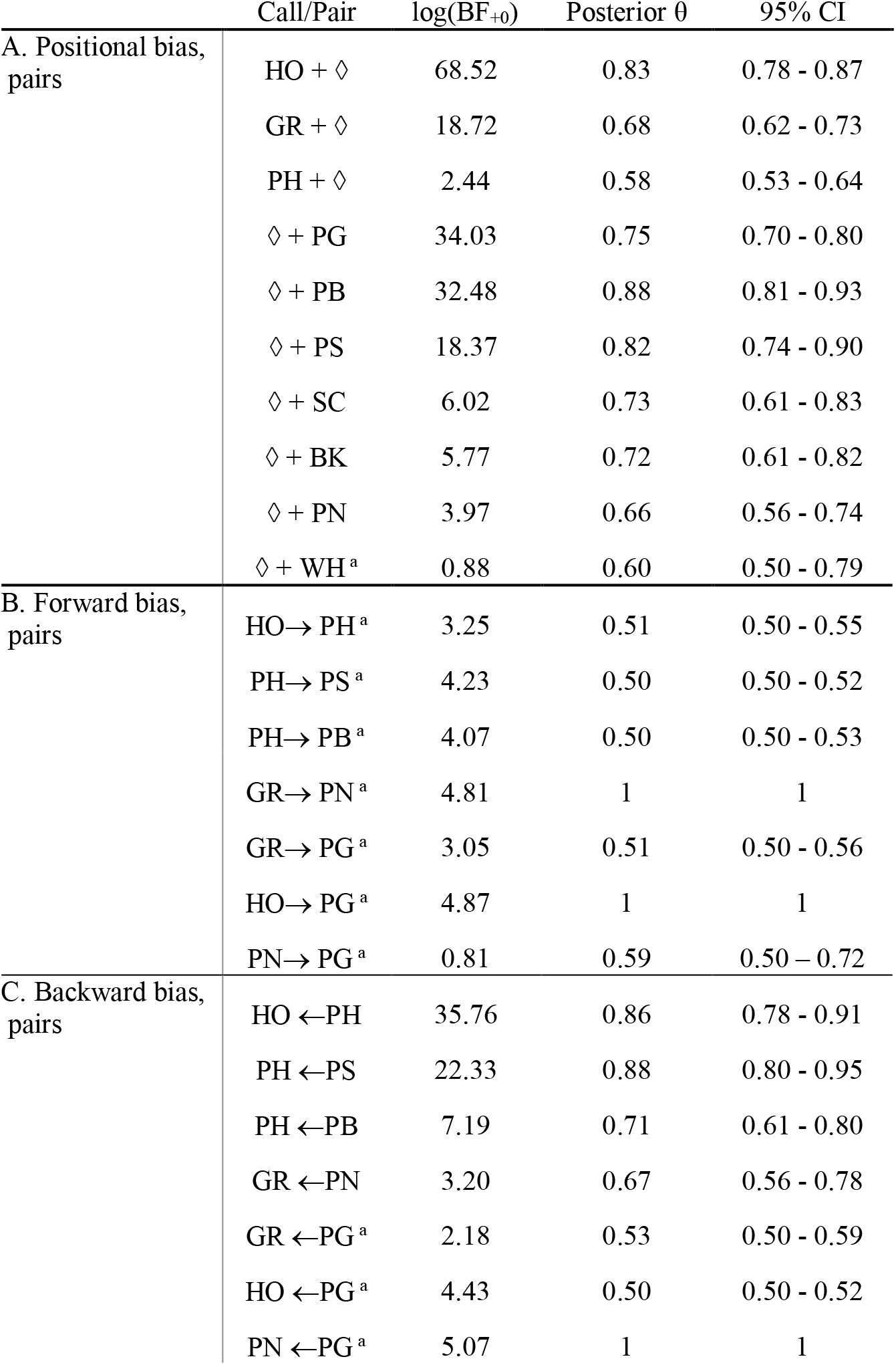
Bayesian binomial test for positional, forward, and backward bias in two-vocal units (pairs).

| Call/Pair | log(BF<sub>+0</sub>) | Posterior $\theta$ | 95% CI |
| --- | --- | --- | --- |
| **A. Positional bias, pairs** |  |  |  |
| HO + $\diamond$ | 68.52 | 0.83 | 0.78 - 0.87 |
| GR + $\diamond$ | 18.72 | 0.68 | 0.62 - 0.73 |
| PH + $\diamond$ | 2.44 | 0.58 | 0.53 - 0.64 |
| $\diamond$ + PG | 34.03 | 0.75 | 0.70 - 0.80 |
| $\diamond$ + PB | 32.48 | 0.88 | 0.81 - 0.93 |
| $\diamond$ + PS | 18.37 | 0.82 | 0.74 - 0.90 |
| $\diamond$ + SC | 6.02 | 0.73 | 0.61 - 0.83 |
| $\diamond$ + BK | 5.77 | 0.72 | 0.61 - 0.82 |
| $\diamond$ + PN | 3.97 | 0.66 | 0.56 - 0.74 |
| $\diamond$ + WH<sup>a</sup> | 0.88 | 0.60 | 0.50 - 0.79 |
| **B. Forward bias, pairs** |  |  |  |
| HO→ PH<sup>a</sup> | 3.25 | 0.51 | 0.50 - 0.55 |
| PH→ PS<sup>a</sup> | 4.23 | 0.50 | 0.50 - 0.52 |
| PH→ PB<sup>a</sup> | 4.07 | 0.50 | 0.50 - 0.53 |
| GR→ PN<sup>a</sup> | 4.81 | 1 | 1 |
| GR→ PG<sup>a</sup> | 3.05 | 0.51 | 0.50 - 0.56 |
| HO→ PG<sup>a</sup> | 4.87 | 1 | 1 |
| PN→ PG<sup>a</sup> | 0.81 | 0.59 | 0.50 - 0.72 |
| **C. Backward bias, pairs** |  |  |  |
| HO ← PH | 35.76 | 0.86 | 0.78 - 0.91 |
| PH ← PS | 22.33 | 0.88 | 0.80 - 0.95 |
| PH ← PB | 7.19 | 0.71 | 0.61 - 0.80 |
| GR ← PN | 3.20 | 0.67 | 0.56 - 0.78 |
| GR ← PG<sup>a</sup> | 2.18 | 0.53 | 0.50 - 0.59 |
| HO ← PG<sup>a</sup> | 4.43 | 0.50 | 0.50 - 0.52 |
| PN ← PG<sup>a</sup> | 5.07 | 1 | 1 |

**Figure 3:**
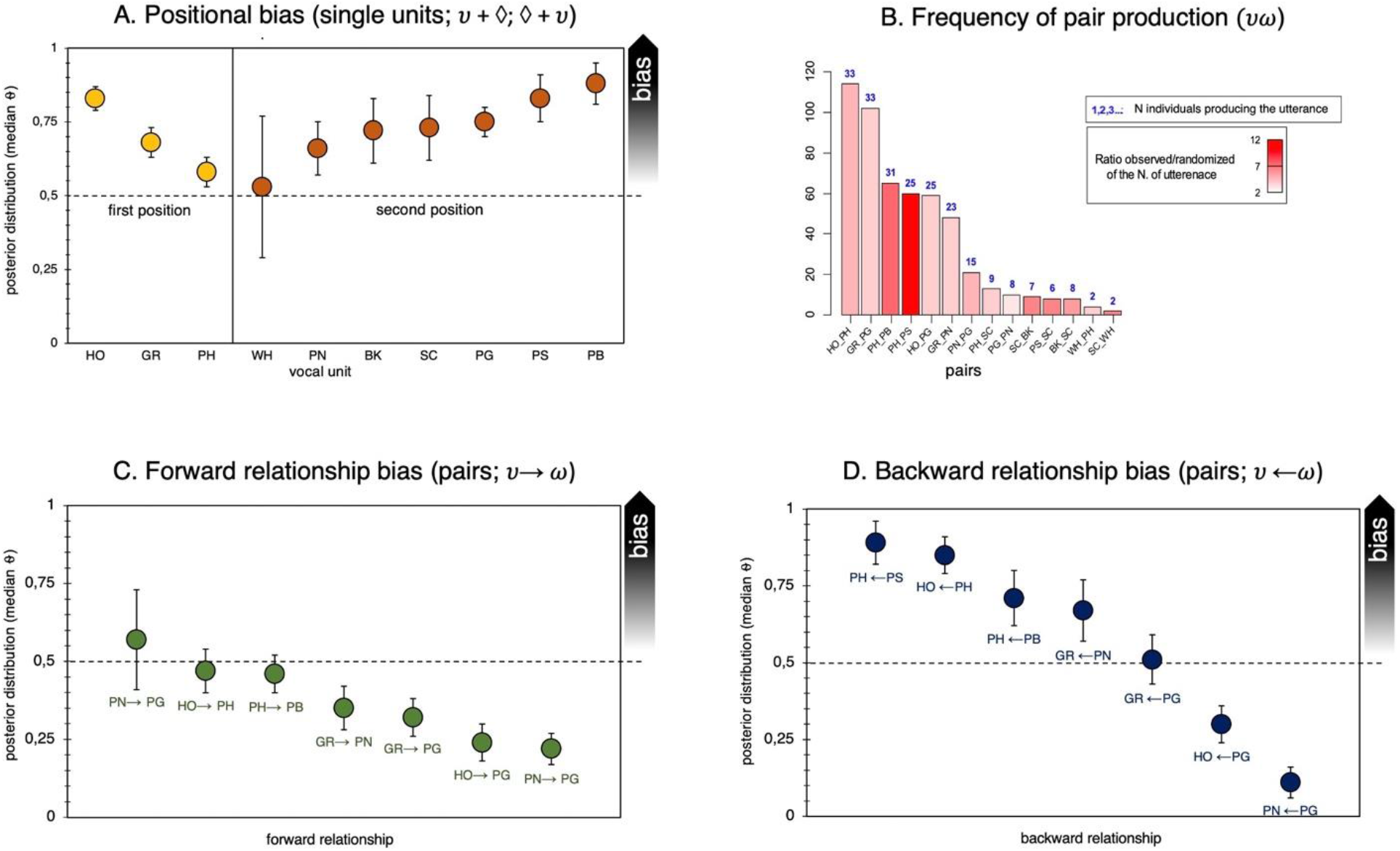
Positional bias, specific ordering frequency, and relationship bias for two-unit vocal sequences (pairs). (A) Likelihood for a certain unit *ν* to occur in first position (yellow) or in second position (orange) in two-unit utterances, expressed as median posterior distribution (θ). Bars stand for Credible Intervals (CI) at 95%. ◊ = irrespective of call type. (B) Frequency of production of the pairs in utterances with two vocal units that were produced above chance (i.e., >95% more likely than by random juxtaposition of single vocal units). The height of each bar corresponds to the number of times each utterance was recorded. The colour gradient in the bars depicts the number of times each utterance was observed divided by the number of times each utterance was present on average in each randomization (averaged over 1000 randomizations). The colour ranges from the lowest ratio in white (i.e., the utterance was present in the observed data only two times more than in the randomization) to the highest ratio in red (i.e. the utterance was present in the observed data 12 times more than in the randomized data). The number on top of each bar in blue indicates the number of individuals that produced each utterance. (C) Likelihood for a certain unit (*ν*) in first position to be in a forward relationship (green) with a certain unit (*ω*) in second position. (D) Likelihood for a certain unit (*ω*) in second position to be in a feedback relationship (blue) with a certain unit (*ν*) in first position, in two-unit utterances.

#### Ordering Frequency

We used a classic randomization procedure based on 1000 randomizations and we found that out of the 58 unique pairs produced by the chimpanzees, 14 pairs in sequences with two vocal units were produced above chance level (i.e., more often than by random juxtaposition of single vocal unit from the repertoire, **Fig. 3B**). Out of the 14 “above chance” pairs, seven were produced by at least 10 individuals (GR_PG, GR_PN, HO_PG, HO_PH, PH_PB, PH_PS, PN_PG).

#### Transitional Bias

We evaluated two different kinds of transitional relationships between the two vocal units forming the pair, thus mimicking information-theoretic concepts used for quantifying processing efforts in the psycholinguistic literature^46–48^. First, we tested how likely it was to find a call *ω* in second position, compared to all other possible follow-on units, given a call *ν* in first position. We called this transitional possibility a “forward relationship”. Second, we tested how likely it was to find a call *ν* in first position, compared to all the possible preceding units, given a call *ω* in second position. We called this alternative transitional possibility a “backward relationship”. None of the seven pairs found above chance showed any forward relationship between the two vocal units within the sequence. Indeed, there was no call in first position that preferentially occurred with a certain call in second position (**Fig. 3C** **and Table S1-S2**). Thus, the first vocal unit in a pair set no constraint on the following vocal unit. The second position did however appear to be bound to certain units in first position. For example, the pair PH_PS occurred 60 times, but while PH was followed by PS 60 times out of 186 pairs, PS followed PH 60 times out of 67 pairs.

Results from the Bayesian binomial test evaluating positional (A), forward (B), and backward (C) bias in two-vocal units (pairs). For each analysis, we conducted Bayesian binomial tests on JASP^49^ with default effect size priors and flat Beta (1,1) to quantify the relative likelihood for a potential positional or relationship bias within the sequence (see Methods). The number of successes over all trials for each unit was defined according to the number of occurrences found in the most frequent position (first or second position) for the positional analysis (A). The number of successes over all trials for each sequence was defined according to the number of times that *ν* was followed by *ω* (or that *ω* was preceded by *ν*), compared to the number of times that *ν* was followed by any other unit (or that *ω* was preceded by any other unit) for the two relationship analyses (B-C). Results are reported for the Bayesian factor B_+_0 (unless indicated otherwise), testing the hypothesis that the proportion of occurrences is higher than the default test value set at 0.5 (50%). Effect size estimates are reported as median posterior population (θ) with Credible Intervals (CI) set at 95% (^50^). ^a^ = Bayesian factor B0+ testing the null hypothesis that the proportion of occurrences in the most frequent position is not higher than the default test value set at 0.5. ◊ = irrespective of call type.

We indeed found four sequences with a strong backward bias—PH-PS, HO-PH, PH-PB, and GR-PN— suggesting that a certain second unit predominantly occurred when it was preceded by another specific unit (**Fig. 3D** **and** **Table 2C**). Regardless of the position of occurrence within the pair, we finally found that when inserting all calls in a bidirectional combinatorial network for the two-unit pairing system, PH was the call that was used the most with all other calls in the dataset (Betweenness Centrality (BC) = 12.45). This suggests that PH might be a key call in the two-unit network that can be emitted flexibly in sequence with all other calls (HO=4.566, SC=4.116, GR=3.816, BK=3.5, PB=2.533, PG=1.115, WH=1.033, PS=0.666, PN=0.2; **Fig. 2B-C**).

### Vocal sequence analysis: three vocal units

The chimpanzees produced 104 unique sequences with exactly three vocal units (**Table S1**). Here we begin to explore how chimpanzees combine simple two-unit sequences with a third unit to produce three-unit sequences. We first assessed which three-unit sequences are produced above chance level and which pairs would appear more than by chance within three-unit sequences. We thus first asked: (1) Specific ordering frequency: a) Do three-unit sequences show fixed order frequencies that go beyond random juxtaposition of single vocal units? And b) Is the position of some pairs occurring within three-unit sequences non-random? (2) Positional bias: Are some pairs biased towards a certain specific position within three-unit sequences (head position—i.e., first and second position bias with a third call taking the final position; tail position—i.e., second and third position bias with a third call taking the head position) within bigger three-unit sequences? (3) Transitional bias: Do relationships between these pairs and single calls exist (i.e., *νω* follows ε, or *νω* precedes ε) when forming three-unit sequences?

(1) *Ordering frequency*. As for the pairs above, we used classic randomization procedures and found that 49 unique sequences with three vocal units were produced above chance level (**Fig. S5**). Out of these 49 above chance sequences, eight were produced by at least 10 individuals (GR_PG_GR, GR_PG_PN, HO_PH_HO, HO_PH_PB, HO_PH_PS, PH_PB_PH, PH_PB_PS, PH_PS_PB, **Fig. 2B**). Interestingly, four of the seven pairs that were produced above chance level and by at least 10 chimpanzees in two-unit sequences (GR_PG, HO_PH, PH_PB, PH_PS) are also found in these eight sequences with three vocal units. When specifically considering all the pairs occurring in three-unit sequences we identified 64 unique pairs. Using the same randomization procedure as for the pairs in sequences with two vocal units we found that 21 out of the 64 unique pairs in sequences with three vocal units were produced above chance level (**Fig. 4A**). Out of these 21 above chance pairs, 13 were produced by at least 10 individuals (GR_PG, HO_PH, PB_BK, PB_PH, PB_PS, PG_GR, PG_PB, PG_PN, PH_PB, PH_PS, PS_PB, PS_SC, SC_PS). Altogether, this suggests that some vocal units are added before or after pairs to produce longer sequences. A comparison of the frequency distribution of pairs in two-unit sequences with the frequency distribution of pairs in three-unit sequences reveals similar frequencies for these four pairs (**Fig. S6**).
(2) *Positional Bias*. We used Bayesian binomial tests (see Methods and **Table 3A**) to assess whether any of the four pairs found above chance at both two- and at three-unit level sequences (GR_PG, HO_PH, PH_PB, PH_PS) showed bias to occur at head or tail position in the three-unit sequences. We found that HO_PH, GR_PG, and PH_PB showed a strong positional bias towards the head position, while no effect was found for PH_PS (**Fig. 4B** **and** **Table 3A** **and S3**).
(3) *Transitional Bias*. We tested for possible relationships between pairs and single units in three-unit sequences (i.e.,[*νω*]ε or ε[*νω*]). First, we asked in one case how likely is it to find ε, compared to all possible following units (“forward relationship”), given *νω* in head position. Second, we asked how likely is it to find ε, compared to all the other possible preceding units (“backward relationship”), given *νω* tail position. Transitional relationships between pairs and single units revealed one strong forward relationship between GR_PG and the following GR unit (**Fig. 4C and Table 3B**), while the two-unit PH_PS was predominantly preceded by the HO unit (**Fig. 4D**, **Table 3C**; **Fig. 5 and Table S4-5**).

**Figure 4:**
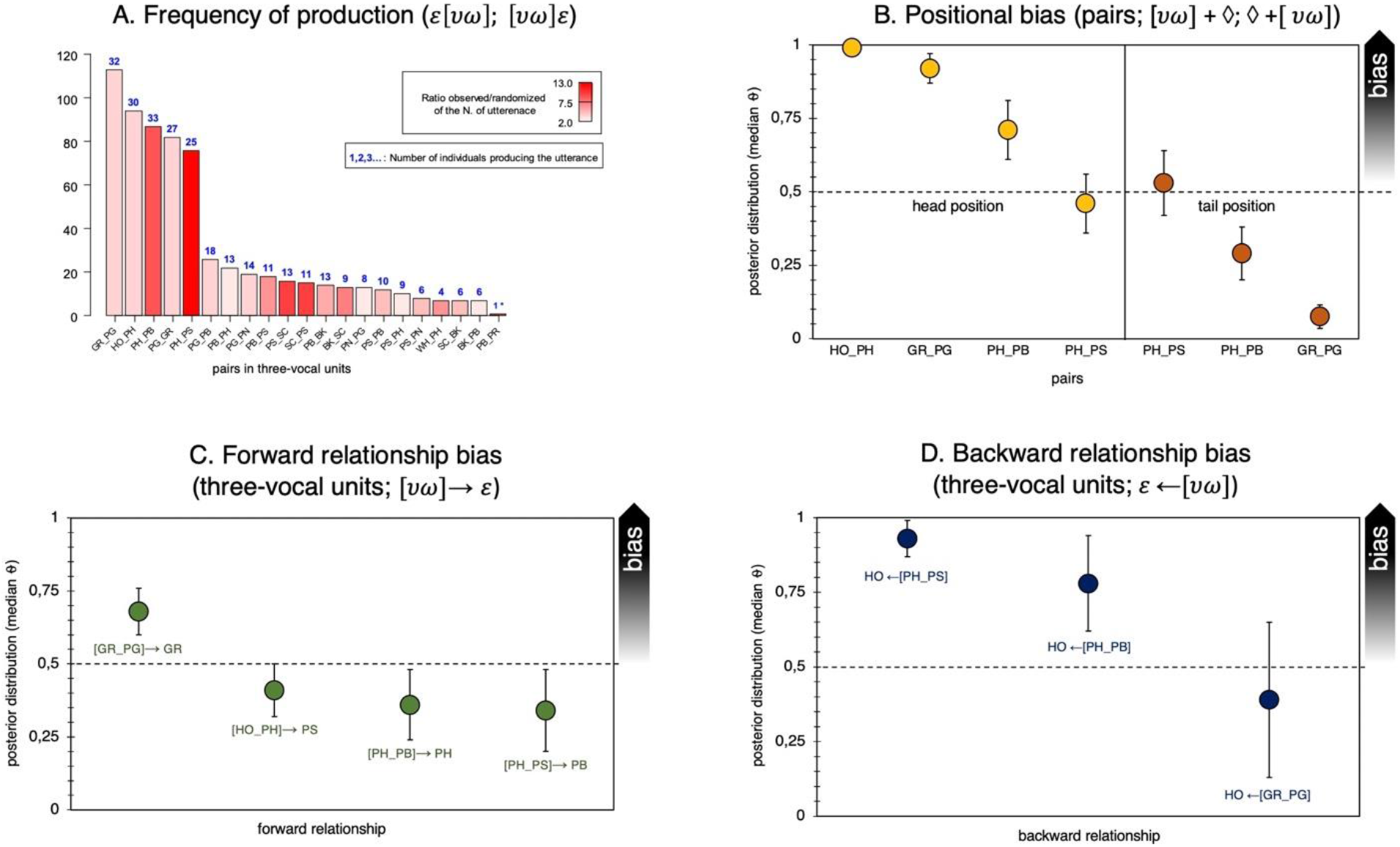
Specific ordering frequency, positional bias, and relationship bias for three-unit vocal sequences. (A) Pairs in utterances with three or more vocal units that were produced above chance (i.e., >95% more likely than by random juxtaposition of single vocal units). The height of each bar corresponds to the number of times each utterance was recorded. The colour gradient in the bars depicts the number of times each utterance was observed divided by the number of times each utterance was present on average in each randomization (averaged over 1000 randomizations). The colour ranges from the lowest ratio in white (i.e. the utterance was present in the observed data only two times more than in the randomization) to the highest ratio in red (i.e. the utterance was present in the observed data 53 times more than in the randomized data). The number on top of each bar in blue indicates the number of individuals that produced each utterance. (B) Likelihood for a certain pair *νω* to occur in head position (yellow) or in tail position (red) in three-unit utterances, expressed as median posterior distribution (θ). Bars stand for Credible Intervals (CI) at 95%. ◊ = irrespective of call type. (C) Likelihood for a certain two-unit utterance (*νω*) to be in a forward relationship (green) with a certain (*ε*) unit following it. (D) Likelihood for a certain two-unit utterance (*νω*) to be in a backward relationship (blue) with a certain (*ε*) unit following it. * The pair PB_PR appeared in the dataset only 1 time and it appeared 250 times less in the randomizations as compared to the observed frequency (and not 13 times as indicated by the red color). We used a gradient of color from 2 to 13 to visualize the variation in the other pairs bearing in mind that the ratio observed/randomized or the last pair (PB_PR) is much higher than indicated in the figure.

**Table 3.** Bayesian binomial test for positional, forward, and backward bias in three-vocal units.

| | Call/Pair | log(BF<sub>+0</sub>) | Posterior $\theta$ | 95% CI |
| --- | --- | --- | --- | --- |
| A. Positional bias,<br>pairs in three<br>vocal units | [HO + PH] + $\diamond$ | 61.23 | 0.99 | 0.96 - 1 |
| | [GR + PG] + $\diamond$ | 47.32 | 0.92 | 0.87 - 0.96 |
| | [PH + PB] + $\diamond$ | 6.70 | 0.71 | 0.61 - 0.80 |
| | $\diamond$ + [PH + PS]<sup>a</sup> | 2.27 | 0.53 | 0.50 - 0.61 |
| B. Forward bias,<br>pairs in three<br>vocal units | [GR + PG] → GR | 5.93 | 0.68 | 0.59 - 0.77 |
|  | [HO_PH] → PS<sup>a</sup> | 3.08 | 0.51 | 0.50 - 0.57 |
|  | [PH_PB] → PH<sup>a</sup> | 3.03 | 0.52 | 0.50 - 0.57 |
|  | [PH_PS] → PB<sup>a</sup> | 2.66 | 0.52 | 0.50 - 0.60 |
| C. Backward bias,<br>pairs in three<br>vocal units | HO ← [PH_PS] | 18.04 | 0.93 | 0.83 - 0.98 |
|  | HO ← [PH_PB] | 3.88 | 0.78 | 0.61 - 0.91 |
|  | HO ← [GR_PG]<sup>a</sup> | 1.35 | 0.58 | 0.50 - 0.77 |

**Figure 5.**
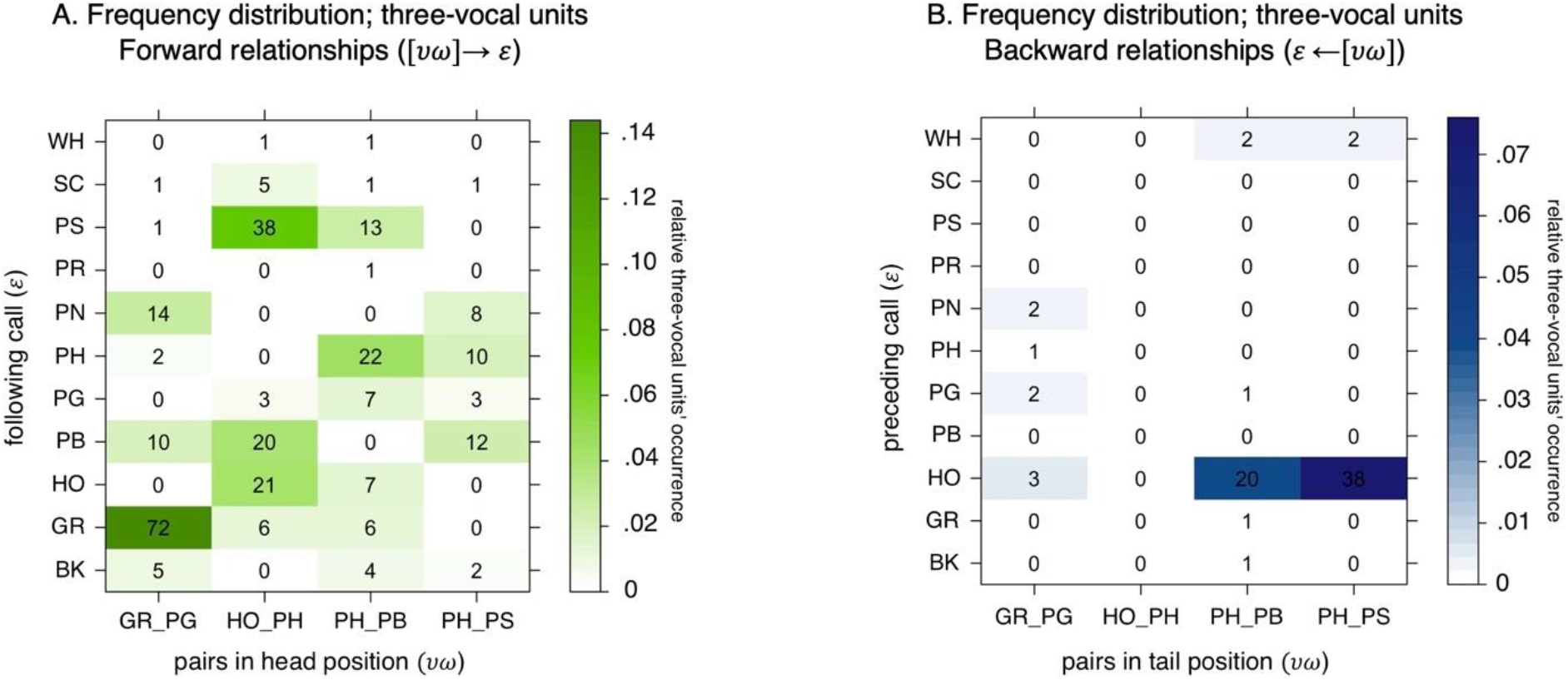
Frequency distribution for three-vocal unit sequences. (A) Frequency distribution for three-vocal unit sequences with two-vocal unit sequences occurring in head position (*νω*) along the x-axis and following call units (*ε*) along the y-axis. Color gradients (white-to-green) represent the relative occurrence of each pair within the three-vocal units’ set. In each cell, the absolute frequency count for each three-vocal unit is also reported. (B) Frequency distribution for three-vocal unit sequences with two-vocal unit sequences occurring in tail position (*νω*) along the x-axis and preceding call units (*ε*) along the y-axis. Color gradients (white-to-blue) represent the relative occurrence of each pair within the three-vocal units’ set. In each cell, the absolute frequency count for each three-vocal unit is also reported.

Results from the Bayesian binomial test evaluating positional (A), forward (B), and backward (C) bias in three-vocal units. The same parameters as described in Table 2 were used (see Methods). The number of successes over all trials for each pair was defined according to the number of occurrences found in the most frequent position (head vs. tail) in the three-vocal units for the positional analysis (A). The number of successes over all trials for each utterance was thus defined according to the number of times that *νω* was followed by ε (or that *νω* was preceded by ε), compared to the number of times that *νω* was followed by any other units (or that *νω* was preceded by any other unit) for the relationship analyses (B-C). Results are reported for the Bayesian factor B_+_0 (unless indicated otherwise), testing the hypothesis that the proportion of occurrences is higher than the default test value set at 0.5 (50%). Effect size estimates are reported as median posterior population (θ) with Credible Intervals (CI) set at 95% (^50^). ^a^ = Bayesian factor B0_+_ testing the null hypothesis that the proportion of occurrences in the most frequent position is not higher than the default test value set at 0.5. ◊ = irrespective of call type.

## Discussion

Compared to human language, in other species, sequences longer than two units that contain meaning-bearing units have not yet been found, or have only been anecdotally described^26,28,29^ (**Fig. 6**). Here we show potential for more complex patterning in the vocal output of chimpanzees, one of our closest living relatives.

**Figure 6:**
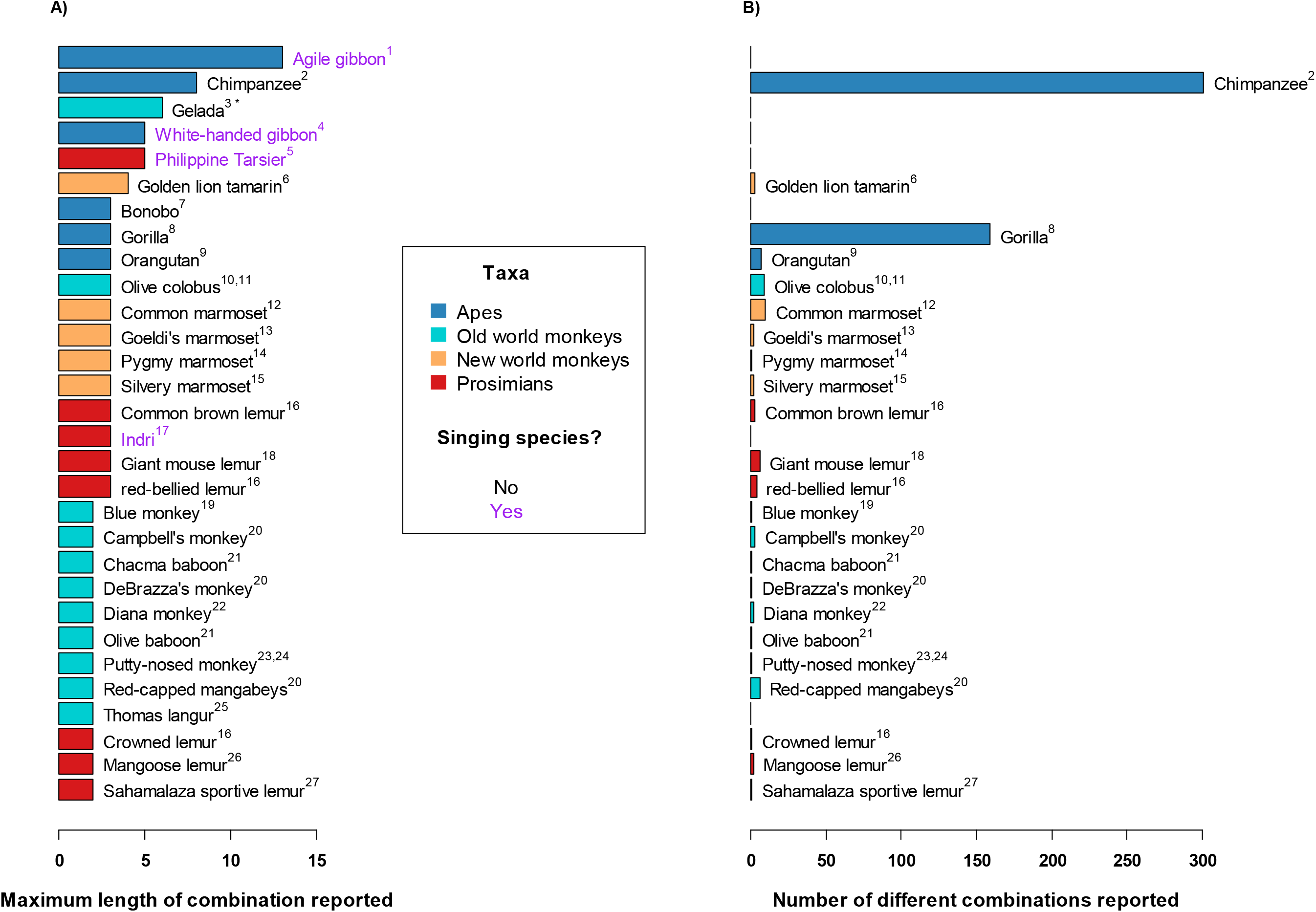
A comparison of vocal sequence production across primates: maximum sequence length and number of sequences. The barplot on the left indicates the maximum length of sequences reported in each species (i.e., the number of unique vocal units emitted in a single sequence) (A). The barplot on the right indicates the number of different sequences reported to comprise at least two different vocal units (B). We counted sequences in which different vocal units occurred at least once. If the same vocal unit was repeated later in the sequence this was not counted as a new sequence. Species showing no number of different sequences may reflect lack of published information. The colour of the bars in both plots indicate the taxa of the species. Apes (including great and lesser apes), old-world monkeys, new world monkeys, and prosimians are depicted in dark blue, light blue, orange, and red respectively. The singing species are indicated by purple text. Singing species are species such as Indris or gibbons which emit different vocal units in song. These units are not necessarily emitted singularly nor in other parts of the vocal repertoire. Superscript numbers refer to the relevant citations listed below. <u>*Notes:*</u> We do not consider repeats of the same vocal unit (e.g., A_A_A) as a sequence. For the sake of comparison with other species in which the methods differ slightly from our current approach, we recalculated for this Figure the length and diversity of sequences excluding counts of repetitions of the same vocal unit at any time point within a sequence (e.g., we treated A_B_D as the same sequence as A_B_A_D). Using Google Scholar, we have tried to conduct an exhaustive literature search of several vocal sequence production across primate taxa. We have only included species in which at least one vocal sequence was clearly reported either in peer-reviewed texts or shown on spectrograms. We cannot exclude that we missed some publications that reported sequences in species not listed here. We also cannot exclude that the species indicated in this figure may emit longer vocal sequences or have a larger variety of sequences than is currently reported in the literature. For several species, it is not clear if the vocal units emitted in the sequences are also produced singularly or if they are only produced as part of a sequence. * Gelada were reported to produce utterances containing up to 26 vocal units in a ‘song-like’ fashion^51^. They possess however only 6 different vocal units that they repeat within the same sequence. Furthermore, the criteria to define which calls are produced during the utterance differs from our study (5-sec intervals maximum between calls versus 1 sec in our study). References: (1)^52^, (2) our study, (3)^51^, (4)^33^, (5)^53^, (6)^34^, (7)^54^, (8)^55^, (9)^56^, (10)^57^, (11)^38^, (12)^37^, (13)^58^, (14)^59^, (15)^60^, (16)^61^, (17)^62^, (18)^63^, (19)^64^, (20)^65^, (21)^66^, (22)^67^, (23)^68^, (24)^69^, (25)^70^, (26)^71^, (27)^72^.

First, we show that chimpanzees have a highly flexible vocal sequencing system, to an extent not yet demonstrated in a non-human primate (**Fig. 6**). Taї chimpanzees produced 390 unique sequences comprising two or more vocal units (**Fig. 1**, **Table S1**). More than one-third of their vocal output includes at least two units, with 15% of vocal sequences containing three to ten units. Note these numbers are likely an underestimation of the vocal sequence potential of chimpanzees since the number of new sequences found had not reached asymptote after nearly 5000 recordings (**Fig. S7**). Our results also illustrate the high degree of flexibility in chimpanzee vocal sequence output since, except for pants, each of the single vocal units occurred in numerous different sequences with at least four different types of vocal units (**Fig. 2B**).

Second, within the vocal flexibility of the chimpanzee, we found evidence of positional and transitional bias using very conservative measures. At the two-vocal unit level, first we found a strong positional bias such that almost each of the single vocal units occur in either the first or second position (e.g. HO most predominantly occurs in first position whereas PB predominantly occurs in last position). Second, there was over-representation of some specific ordered pairs. Third, we found a strong transitional bias within the overrepresented pairs, such that the call in second position was predominantly emitted with a specific call in first position (e.g., PH was preceded predominantly by HO). We conducted the same analyses at the three-vocal unit level. Our fourth finding showed over-representation of about a third of the three-unit sequences. Fifth, we found that the same pairs showed a specific order in both the two- and three-vocal units’ sample (e.g., HO_PH occurs as a pair but can also occur with an added third unit: HO_PH_PB or HO-PH_PS). Sixth, we found strong positional bias, such that these over-represented pairs occurred in either head or tail positions (e.g., third units added to HO_PH typically occur after and not before the pair is emitted: HO_PH_X). Finally, we found a strong transitional bias within the overrepresented pairs, such that some pairs in head/tail position were most likely emitted with a certain other call in tail/head position. For example, the pair GR_PG was more likely to be followed by GR than by any other singe vocal units while the pair PH_PB was more likely to be preceded by HO than by any other vocal unit).

Overall, the data suggest that chimpanzees might possess some fundamental combinatorial principles which can be used to establish sequential relationships across multiple sound units. Positional information in two-unit sequences is relevant to the sequential ordering of the uttered message and most importantly, the first position seems to be filled with compulsory information when a certain call needs to be introduced in the second position. This seems to be especially relevant for calls like PS or PB, which occur less frequently in isolation and which are almost always preceded by the specific call PH in two-unit sequences. The fact that chimpanzees have access to local ordering patterns and precedence relationships during spontaneous vocal production supports previous data coming from auditory sound discrimination paradigms in other primates. The latter demonstrate sensitivity to ordering violations between adjacent sound units in artificial strings generated by simple finite-state grammars of the (AB)^2^ type^13,16,73–75^. Beyond adjacency relationships, chimpanzees and other primates have been shown to generalize over dependency rules between non-adjacent elements in both sound and visual discrimination tasks^15,17,76,77^. This suggests some ordering capacity that would go beyond simply adding two vocal units together in a sequence. In this study, we found that at the three-unit level, chimpanzees might also rely on positional information, where some over-represented pairs within the sequence are consistently produced at the beginning of the sequence (first position, e.g., HO_PH, GR_PG, PH_PB). Similarly, a few over-represented pairs—which occur above chance at both two-unit and three-unit level—appear to co-occur together with a third vocal unit, which generally linearly precedes or follows them, when occurring in three-unit sequences (e.g., GR_PG + GR, HO + PH_PS, HO + PH_PB).

Our results also indicate that some two-unit sequences are reused as ready-combined calls in three-unit sequences. This might offer intriguing evidence for a potential pairing prerequisite during vocal communication in animals^26,27^. This would mean that chimpanzees might form combined meanings from two individual calls, which can, in turn, be recombined with a third call to eventually output a third combined meaning. However, without looking into contextual information, which goes beyond the scope of the present study, our analysis cannot exclude that the pairing effects in the three-vocal units result from simple transitional relationships between adjacent elements. In this respect, three-unit sequences with over-represented pairs, like HO←[PH-PS] or HO←[PH-PB], might be rather analysed as HO←PH←PS or HO←PH←PB, where vocal units in position two and three are strongly dependent on the vocal unit occurring in the preceding position. This would suggest that linear order in chimpanzees would suffice to create longer sequences, notwithstanding transitional relationships between the internal elements. Worth noting, the fact that most two-unit pairs can be produced with either unit emitted first, and that pairs can also be preceded or followed by other units (**Fig. 5** **and S5**) suggests that biologically-induced auto-correlation effects cannot fully explain the observed patterns. For example, several pairs found in utterances with three vocal units can be preceded and/or followed by four to eight different single vocal units (**Table S4**, **Fig. 5)**. Likewise, within pairs, although we find ordering effects, we find occurrences of most pairs being emitted with either element first (**Fig. 2A** **and** **2B**). It is thus unlikely that the sequences emitted by chimpanzees are purely the product of anatomical constraints requiring certain vocal units to be produced before others.

Sequences longer than two units included more than 60 different sequences, together representing 15.8% of all utterances emitted. One interpretation might be that longer sequences are not important when emitted so infrequently. Alternatively, one can argue that frequency does not necessarily equate to significance, as rare but crucial parts of vocal repertoires are well documented. Alarm calls in social species, for example, are likely underrepresented in overall vocal emission compared to social spacing calls, such as contact calls^65^.

Although the current dataset is not small, determining the importance of these vocal sequences in chimpanzees’ communicative needs will require even larger datasets. A future goal is to determine whether the flexibility we have identified across the chimpanzee vocal repertoire relates to flexibility in the information conveyed. This will require analyses testing, first, for potential contextual and/or meaning shifts between vocal units emitted alone, and the same unit emitted paired with another vocal unit. A second aim is to test whether a pair produced alone undergoes a meaning shift when combined with third or fourth vocal units.

Two-unit and longer sequences are emitted by both sexes, they occur throughout the chimpanzee vocal repertoire, and they are not limited to specific sequences, such as loud calls, nor more specifically to the well-described four-unit pant-hoot sequence (**Fig. S3 A, B, and C**)-which includes single hoos (HO) – panted hoos (PH) – panted screams (PS), and let down phase (panted roars (PR) or pants (PN))^41,78^. Whilst our two-unit sequence analysis includes three sets of pairs that describe part of the pant hoot sequence (head: HO_PH, PH_PS; tail: PH_PS), the other pair GR-PG which appears in tail and head position is not part of pant-hoot sequence. Likewise, many other frequent (above chance) two- and three-unit sequences we have identified are not part of the pant hoot sequence (e.g., HO_PG, GR_PG, GR_PG_PN, GR_PG_PB, PG_PB_BK **Fig. 3A**, **S3 D, E and F, and S5**). Our approach thereby proves valuable in identifying sequences with two vocal units or longer that follow non-random ordering of vocal units that go beyond the commonly described HO_PH_PS sequence.

Other primate species also produce vocal sequences with more than two units. These species fall into two main categories: singers and non-singers (**Fig. 6**). As in other taxa with monogamous breeding systems such as birds, monogamous primates demonstrate co-evolution of singing and duetting, likely due to sexual selection pressures. Across primate clades including prosimians, lesser apes, and the harem structured old world monkey, Geladas, singing usually occurs in territorial or courtship contexts^79,80^, with vocal sequences containing 3-13 unit types^52^ (**Fig. 6**). As in other singing taxa, such as birds and whales^32^, it is thought that the pattern of single units in the primate vocal sequences primarily function to identify individuals or groups and has limited impact on the contextual meaning conveyed to others (^51,62,81^). Units used in ‘song’ are often not used singly, nor in other parts of the vocal repertoire^62,82^. Vocal sequences also occur in some non-singing primates other than chimpanzees, but are mainly restricted to long distance calls and range from 3-5 vocal units, such as in common marmosets^37^, tarsiers^53^, and orangutans^56^ (see **Fig. 6** for a larger list of examples).

Whether vocal sequences also occur in other parts of these repertoires may be under described (**Fig. 6**). Across the primate literature, outside of song and long distance calls, we only found evidence of vocal sequences containing more than two unit types in great ape species: e.g., bonobos emit vocal sequences of at least three call types in food contexts, with variations thought to transmit different information about food preference^83^. To determine whether or not this is a great ape - or a broader primate - phenomenon, we advocate a whole-repertoire approach to vocal analyses to determine the extent of flexible vocal sequence production in primates.

Compared to animal communication, human language compositionality is based on hierarchical structure rather than linear order, where the structure is determined by the word categories being combined (e.g., nouns, verbs, prepositions forming noun phrases, verb phrases, or prepositional phrases, respectively). A ubiquitous phenomenon in human language is indeed the fact that the same linear order may convey different meanings, depending on relevant kinds of the underlying structure. For example, the expression *the man drew a boy with a pencil* can either mean that a man used a pencil to draw a boy, or that he drew a boy who’s holding a pencil. This has often been taken as evidence for the fact that linear ordering might not be a sufficient testing tool for examining the evolution of language^6,84,85^.

An influential hypothesis in linguistic theory states that the computational system holding hierarchical representations in human language—i.e. Merge— might be based on a very parsimonious computation, which builds together phrases and sentences from individual word categories and which is assumed/proposed to be neutrally hardwired in the human brain^86–88^. Some studies have raised the prospect of using Merge as way to describe animal vocal constructions at a higher degree of formalization^26,28^. In contrast to language, however, animal communication seems to lack any categorical dimension on the units of analysis as a hierarchical prerequisite^85^. While cross-species comparability can be premature, the positional bias, the transitional relationships, and the potential for a pairing system in the multi-unit vocal sequences suggests that chimpanzee vocal sequences are a valuable opportunity to examine first, whether the vocal units combine into patterns (longer than two) that change meaning depending on the order of the units, and whether such patterns are further recombined to produce additional shifts in meaning.

## Conclusions

Here, we reveal potentials for a highly flexible animal vocal communication system to contain ordering and combinatorial properties. The chimpanzee two-unit and three-unit vocal sequences revealed positioning bias, transitional relationships, and potential for a pairing system. To analyse the longer sequences larger datasets are required. Purely in terms of patterning, chimpanzee vocal sequences present a valuable opportunity to examine whether the detected patterns also relate to predictable meaning shifts, especially as chimpanzee vocalisations have previously demonstrated relatively rich context specificity. Further research is required to address the challenge of comparing the contexts of production of single units (such as a panted bark), a single unit emitted paired with another vocal unit (panted hoo + panted bark), or a pair emitted before or after a third vocal unit (hoo + [panted hoo + panted bark]; or [panted hoo + panted bark] + grunt). Our results suggest that conclusions from other studies, which state that non-human animal vocal sequences of longer than two meaning-bearing units lack ordering properties, may be premature.

## Methods

### Study site and subjects

We conducted the study within the Taï Chimpanzee project^89^ on wild western chimpanzees at the Taï National Park, Cȏte d’Ivoire (5°45′N, 7°07′W). TB collected data on all adult and subadult chimpanzees from three communities (East, North and South) between two study periods: January-February 2019 and December 2019 to March 2020. We defined as adult all chimpanzees ≥15 years of age and subadult as chimpanzees between 10 and 15 years of age. Adult and subadult chimpanzees are referred in our study as mature individuals. For this study TB collected data on 46 mature chimpanzees, 5 males and 10 females in East group, 4 males and 8 females in North group, and 5 males and 14 females in South group.

### Data collection

TB followed the chimpanzees from dawn to dusk during c.a. 12 hours per day. TB recorded vocalisations during half-day focal animal sampling^90^ switching the focal animal around 12:30 pm, resulting in c.a. 6 h of continuous sampling per focal. Using a 2 s pre-record option, she audio recorded each vocalisation from the focal chimpanzee as well as any vocalisation produced by individuals visible around the focal animal for whom the identity of the caller could be identified with certainty *Ad libitum*^90^. TB recorded the vocalisations using a Sennheiser ME67 directional microphone (digitized at a 48 kHz sampling rate and 24-bit sampling depth) connected to a Tascam DR-40X digital recorder. TB focalled mature chimpanzees for 513 hr and collected ad libitum data for an additional 387.8 hr. This resulted in mean ± SE 13.2 ± 0.9 SE hours of focal sampling on 39 mature focal individuals. In addition, vocal production from seven extra mature individuals was recorded ad libitum. Per individual, TB collected an overall mean ± SE of 34.4 ± 3.5 vocal utterances during focal hours and 85.89 ± 7.02 vocal utterances *Ad libitum*. Per hour of focal data, TB obtained 3.9 ± 0.3 vocal utterances, from which 2.1 ± 0.2 where single vocal units and 0.9 ± 0.1 were sequences (i.e., at least two different vocal units produced one after the other with less than 1-sec pause between them). Of the single vocal units, 1.8 ± 0.2 were non-panted single calls (HO; GR; BK; SC; WH; NV) and 0.4 ± 0.1 were panted single calls (PH; PG; PB; PS; PR).

### Construction of Vocal Repertoire

The chimpanzee vocal repertoire consists of several call types. Most of these call types are emitted either singly or in a ‘panted’ form whereby a voiced inhalation is inserted between each call (**Fig. S2a-b**). Thus hoos can be emitted as single hoos, as repetitions of single hoos with 100 −500 ms between each hoo, or as sequences of panted hoos, with c.a. 100 ms between each hoo and pant. Whilst each call type can be emitted singly, panted versions are only emitted as repetitions. Given that inserting pants between single calls often changes the context in which calls are emitted, we attribute panted versions as being different units. Thus, we divide the repertoire into call types and their panted versions resulting in 7 single forms and 5 panted forms, here termed ‘units’ (see details in Table 1 and **Fig. S2**). Each of these vocal units can be combined, where different combinations result in different sequences. We defined a <u>single utterance</u> as a unit emitted alone or repeated within 2 s intervals. We defined a <u>sequence</u> as different types of vocal-units were emitted with less than 1 s interval (e.g., hoos followed by grunts; or hoos followed by panted hoos emitted within 1s, **Fig. S3c**). Different variants of the same call type (e.g., ‘rest’ or ‘alert’ hoos in ^91^) are not differentiated here but are considered as the same unit type.

### Assigning units to recorded vocal sequences

Recordings are examined using PRAAT spectrograms which show the frequency distribution across the call^92^. An inherent problem in acoustic recordings in tropical forests is the dense background noise, making automated processes for extracting acoustic measurements problematic, especially for quieter calls such as hoos and grunts. However, different call types can be distinguished using spectrograms, which reveal both temporal and spectral properties^41,93^. Call types can be differentiated due to their distinctive acoustic features (Table 1, see SI for spectrograms & sound files of each call type, and gradations of these call types). For the analysis, we considered only calls of high quality, with the lowest frequency band visible, recorded from the beginning to the end, and with the signaller ID defined.

The chimpanzee vocal repertoire is a graded system (**Fig. S8**), such that most call types grade into other call types. Call types that were thus difficult to categorize were sent to a blind coder and an expert in chimpanzee vocal repertoire (CC). If there was no agreement between at least two coders, the call was categorized as “unclear”. We did not include in the analysis utterances containing unclear calls. Utterances in which the start or end could not be coded due to overlap or recording omission were also not included. TB coded all the data. 6% of the data (301 calls across all call types) was subjected to inter-ratter reliability with a blind coder. At the end of the training, TB and the blind coder reached a 94.6 % of agreement on the call classification. In total, 4826 utterances were used for this study out of 5517 utterances recorded, comprising 401 differently constructed utterances which included one to ten different vocal units (see details in Table S1).

### Sequence Analyses

Our goal was to assess whether the sequences formed by two and three vocal units in our study were just a random juxtaposition of single vocal units, or if they conversely resulted from some ordering rules/ non-random order among the individual calls. Sequences formed by three vocal units constituted the maximum vocal sequence length that was reached across all individuals (**Fig. 1**). Thus, the sample sizes of sequences longer than three units were overall too small to run meaningful statistical analysis on each respective length (i.e., four and then five and then six etc.).

#### Two vocal units

Here we asked three questions concerning the patterning organization of single units emitted in sequences of two vocal units (hereafter pairs). (1) Positional bias: Are the vocal units produced by chimpanzees biased towards a certain specific position within a pair? (2) Specific ordering frequency: Do the pairs show fixed order frequencies that go beyond random juxtaposition of single vocal units? (3) Transitional bias: Do relationships between these units exist (i.e., *ν* follows *ω*, or *ν* precedes *ω*) when forming sequences of two vocal units?

(1) *Positional Bias*. A unit *ν* occurring more often in position 1 than in position 2 (or vice versa), would be classified as having a potential positional bias towards position 1 (or vice versa). For each unit, we then conducted a Bayesian binomial test on JASP (^49^) with default effect size priors and flat Beta (1,1) to quantify the relative likelihood for a potential positional bias within the sequence. We defined the number of successes over all trials for each unit according to the number of occurrences found in the most frequent position. Results are reported for the Bayesian factor B+0 (unless reported otherwise) testing the hypothesis that the proportion of occurrences in the most frequent position is higher than the default test value set at 0.5 (50%). Effect size estimates are reported as median posterior population (θ) with credibility intervals (CI) set at 95% (^50^).
(2) *Ordering frequency*. We used randomization routines to assess which sequences comprising two units were produced more than by chance. We first established the frequency of production of each single vocal unit produced singly and in sequences (see **Table 1**). This constituted the frequency pool of observed frequency of call production. We sampled randomly 1000 times as many pairs as the number of pairs recorded for this study and compared each time the distribution of the random pairs to the one of the observed pairs. Pairs were considered to occur more than by chance if the observed frequency was above the randomized frequency in at least 950 randomizations (i.e. 95% of the randomizations).
(3) *Transitional Bias*. We took, for consistency, only the two vocal unit sequences that were produced above chance levels in (2). We tested two different transitional relationships between the two vocal units *ν* and *ω*. First, we asked: given *ν*, how likely is it to find *ω*, compared to all other possible following units? For convenience, we called this transitional possibility a “forward relationship”. Second, we asked: given *ω*, how likely is it to find *ν*, compared to the all possible preceding units? We called this alternative transitional possibility “backward relationship”. We quantified the likelihood for a potential relationship bias between any two units using Bayesian binomial tests with the same parameters and results report as described in (1) above. The number of successes over all trials for each sequence was defined according to the number of times that *ν* was followed by *ω* (or that *ω* was preceded by *ν*), compared to the number of times that *ν* was followed by any other unit (or that *ω* was preceded by any other unit). Since the test value for this Bayesian analysis was kept constant at 0.5, we measured how far above this level a certain call would either precede or follow another call, using a very conservative ratio of combinatorial patterning of 1:1 as a starting point. We thus investigated whether certain calls *ω* exist, which would either precede or follow *ν* at least 50% of the times, compared to all the other calls pooled together.

#### Three vocal units

Here we examined how chimpanzees combine simple two-unit sequences with a third unit to produce longer sequences. We first assessed which three-unit sequences were produced above chance and which pairs of vocal units would appear more than by chance within three-unit sequences. We first asked: (1) Specific ordering frequency: (a) Were the three-unit sequences produced beyond simple random juxtaposition of single vocal units? (b) Were some pairs occurring within three-unit sequences beyond simple random juxtaposition of single vocal units? (2) Positional bias: Are these pairs biased towards a certain specific position (head position, tail position) within bigger three-unit sequences? (3) Transitional bias: Do relationships between these two-unit sequences and single calls exist (i.e., *νω* follows δ, or *νω* precedes δ) when forming three-unit sequences?

(1) *Ordering Frequency*. We repeated the same randomization process as described in “Two-vocal units” above for the three-unit sequences. As for the pairs, the three-unit sequences were considered to occur above chance level if their observed frequency was above the randomized frequency in 950 randomizations. As for (1b) we extracted from each sequence with three vocal units the two possible pairs within these sequences. For instance, a sequence HO_PH_PG would produce the two following pairs: HO_PH, PH_PG (see **Table 1** for the abbreviation of call names). We then assessed which of these pairs in sequences with three vocal units were produced more than by chance using the same approach as for the pairs produced in sequences with only two vocal units. From these analyses, we then moved to question (2).
(2) *Positional Bias*. We restricted our examination to those two-unit sequences which occurred above chance levels in (1b) above. We counted the positional occurrence—head position, i.e., first and second position; tail position, i.e., second and third position—of each two-unit sequence within each of the sequences with three vocal units. Each pair in a sequence was assigned either to head or tail position bias, according to its position of predominant occurrence. Thus, a three-unit sequence *νω*ε that contains a two-unit sequence *νω* occurring more often in head position than in tail position, would be classified as having a potential positional bias towards the head position. For each sequence, we quantified the likelihood for a potential positional bias using Bayesian binomial tests with the same parameters and results report as described in (“Two-vocal units”, 1) above. The number of successes over all trials for each sequence was defined according to the number of two-unit utterances found in the most frequent position.
(3) *Transitional Bias*. We again asked in one case if given *νω*, how likely it is to find ε, compared to all possible following units (“forward relationship”). In the other case, we asked: given *νω*, how likely is it to find ε, compared to all possible preceding units (“backward relationship”)? The potential relationship bias was assessed using frequentist and Bayesian binomial tests with the same parameters and results report as described in (“Two-vocal units”, 1) above. The number of successes over all trials for each utterance was thus defined according to the number of times that *νω* was followed by ε (or that *νω* was preceded by ε), compared to the number of times that *νω* was followed by any other units (or that *νω* was preceded by any other unit). To assess the bias, only the sequences for which ε followed *νω* (or ε preceded *νω*) with a frequency higher than the sum of all other units following *νω* (or preceding *νω*) were included in the analysis.

## Acknowledgements

We thank the Ministère de l’Enseignement Supérieur et de la Recherche Scientifique and the Ministère de Eaux et Fôrets in Côte d’Ivoire, and the Office Ivoirien des Parcs et Réserves for permitting the study. We are grateful to the Centre Suisse de Recherches Scientifiques en Côte d’Ivoire for their logistical support, and to Kayla Kolff and the staff members of the Taï Chimpanzee Project for their support and assistance in collecting the data. This study was funded by the Max Planck Society and the European Research Council (ERC) under the European Union’s Horizon 2020 research and innovation program awarded to C.C. (grant agreement no. 679787). Core funding for the Taï Chimpanzee Project has been provided by the Max Planck Society since 1997. We thank Julia Fischer and Cat Hobaiter for extremely helpful comments on a previous draft of this manuscript.

