## Supplementary Tables and Figures for "Chimpanzees use numerous flexible vocal sequences with more than two vocal units: A step towards language?"

**Supplementary Methods**

Randomization procedure:

We first established the frequency of production of each single vocal unit produced singly and in sequences (see **Table 1**). This constituted the frequency pool of observed frequency of call production.

For the sequence with two vocal units, we sampled randomly from this distribution 817 pairs of single calls, with both calls being different to create 817 random pairs. 817 was the number of sequences with two vocal units recorded for this study (Figure 1). We then compared the distribution of the random pairs to the one of the observed pairs (the pairs that have been recorded) and identified which pairs had an observed frequency above the random frequency. We repeated this process 1000 times to establish whether each pair occurred more than by chance (i.e., more than by random juxtaposition of single vocal unit). Pairs were considered to occur more than by chance if the observed frequency was above the randomized frequency in at least 950 randomizations (i.e., 95% of the randomizations).

For the sequences with three vocal units, we used the same procedure by extracting randomly 458 sequences with three vocal units (i.e., the number of sequences with three vocal units recorded for this study Figure 1) and comparing this distribution to the distribution of observed sequences with three vocal units 1000 times.

For the pairs within sequences with three vocal units, we extracted from each sequence with three vocal units all of the possible pairs within these sequences. For instance, a sequence HO_PH_PG_GR would produce the three following pairs: HO_PH, PH_PG, PG_GR (see **Table 1** for the abbreviation of call names). We then assessed which of these pairs in sequences with three vocal units were produced more than by chance using the same approach as above. For this analysis, we sampled 916 pairs 1000 times.

**Supplementary Figures**

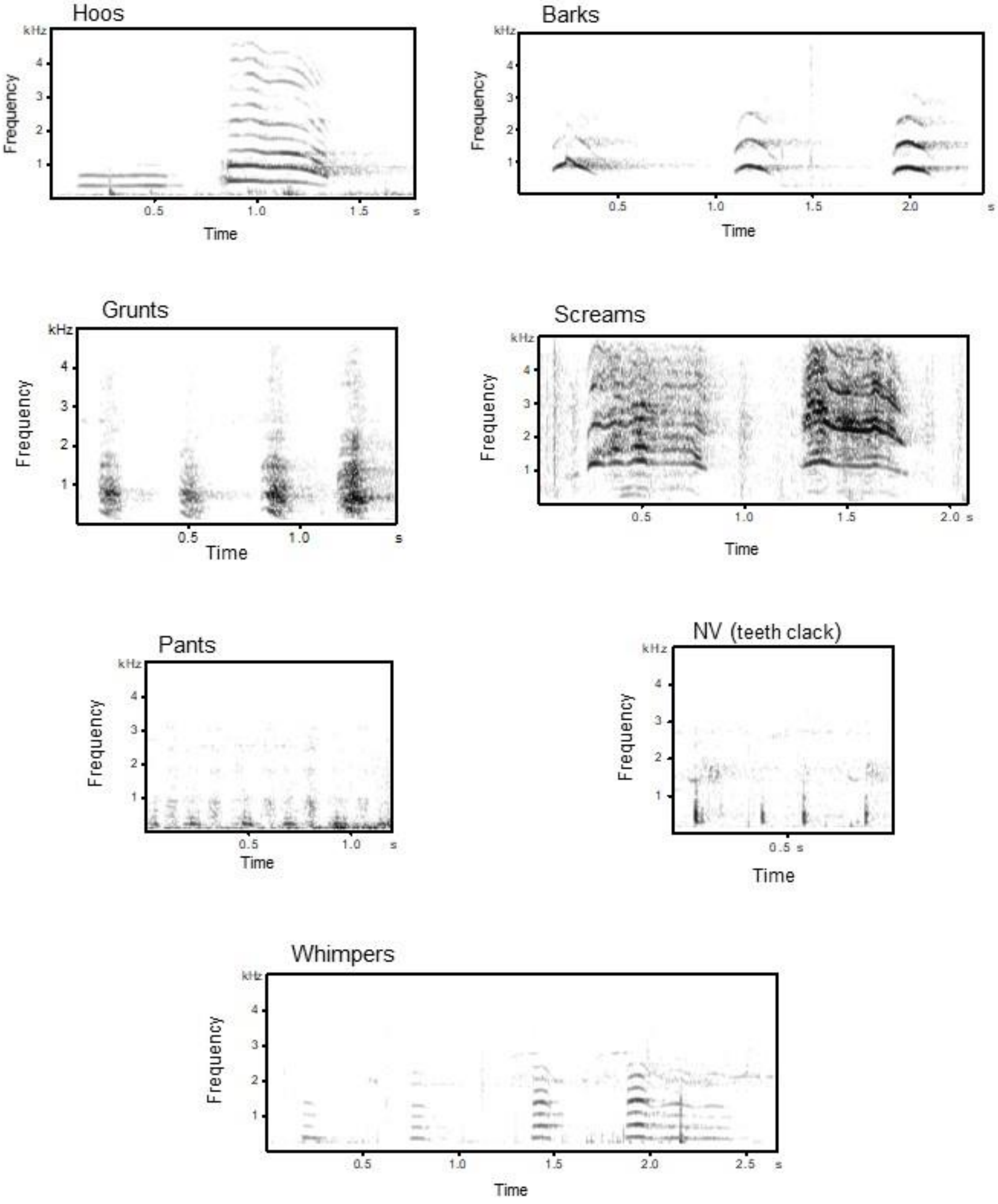


**Figure S1.a:** Spectrograms of chimpanzee single-unit vocalizations.

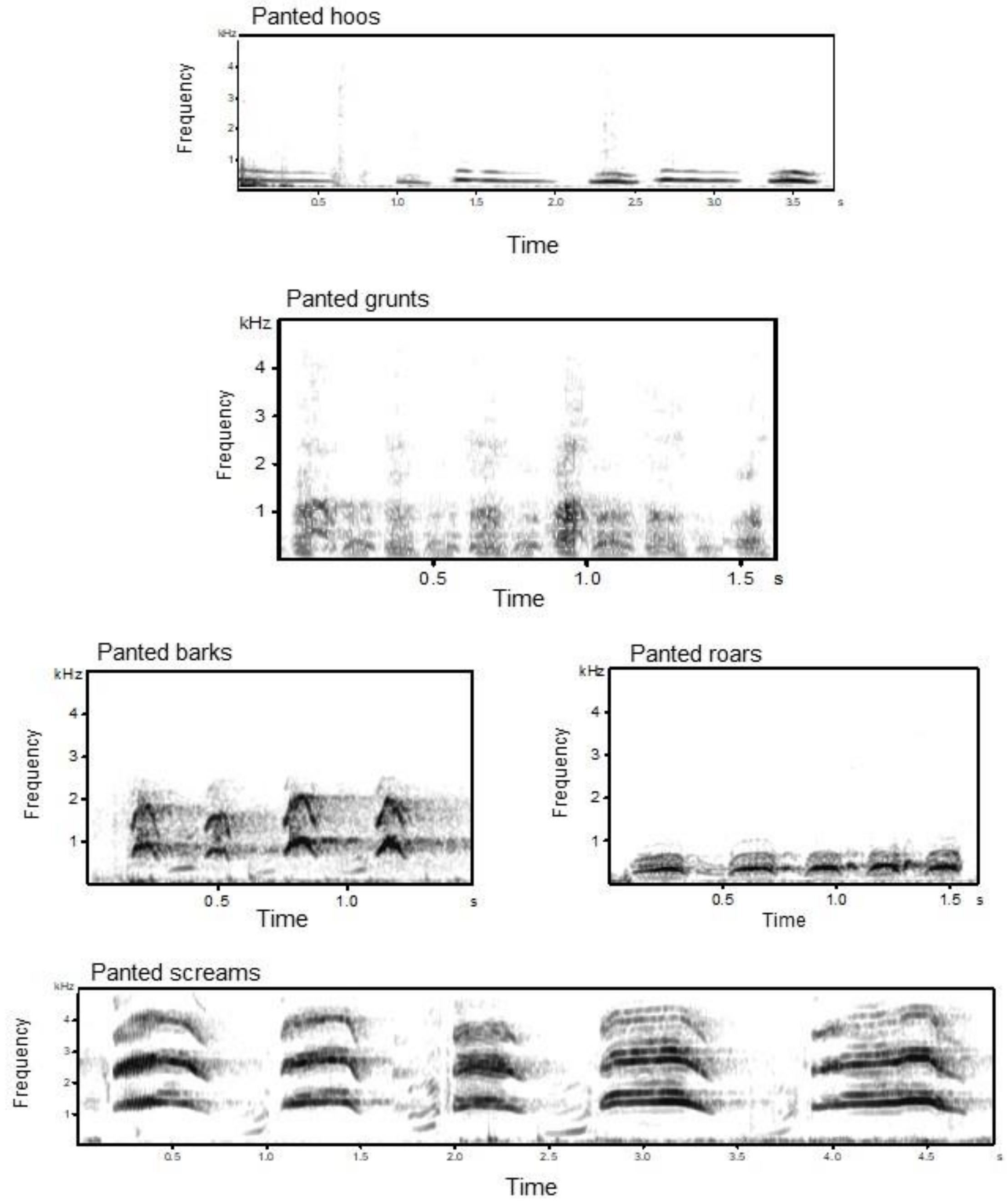


**Figure S1.b:** Spectrograms of chimpanzee panted-unit vocalizations. Roars only occurred as a panted form.

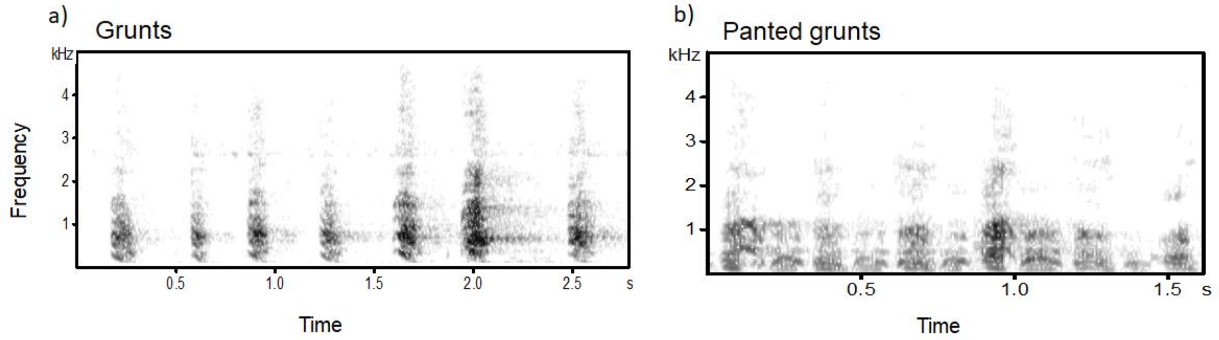


**Figure S2: An example showing the construction of chimpanzee single units.** Spectrograms show (a) a single non-panted call (e.g., series of Grunts) and (b) a single panted call (e.g., series of Panted grunts).

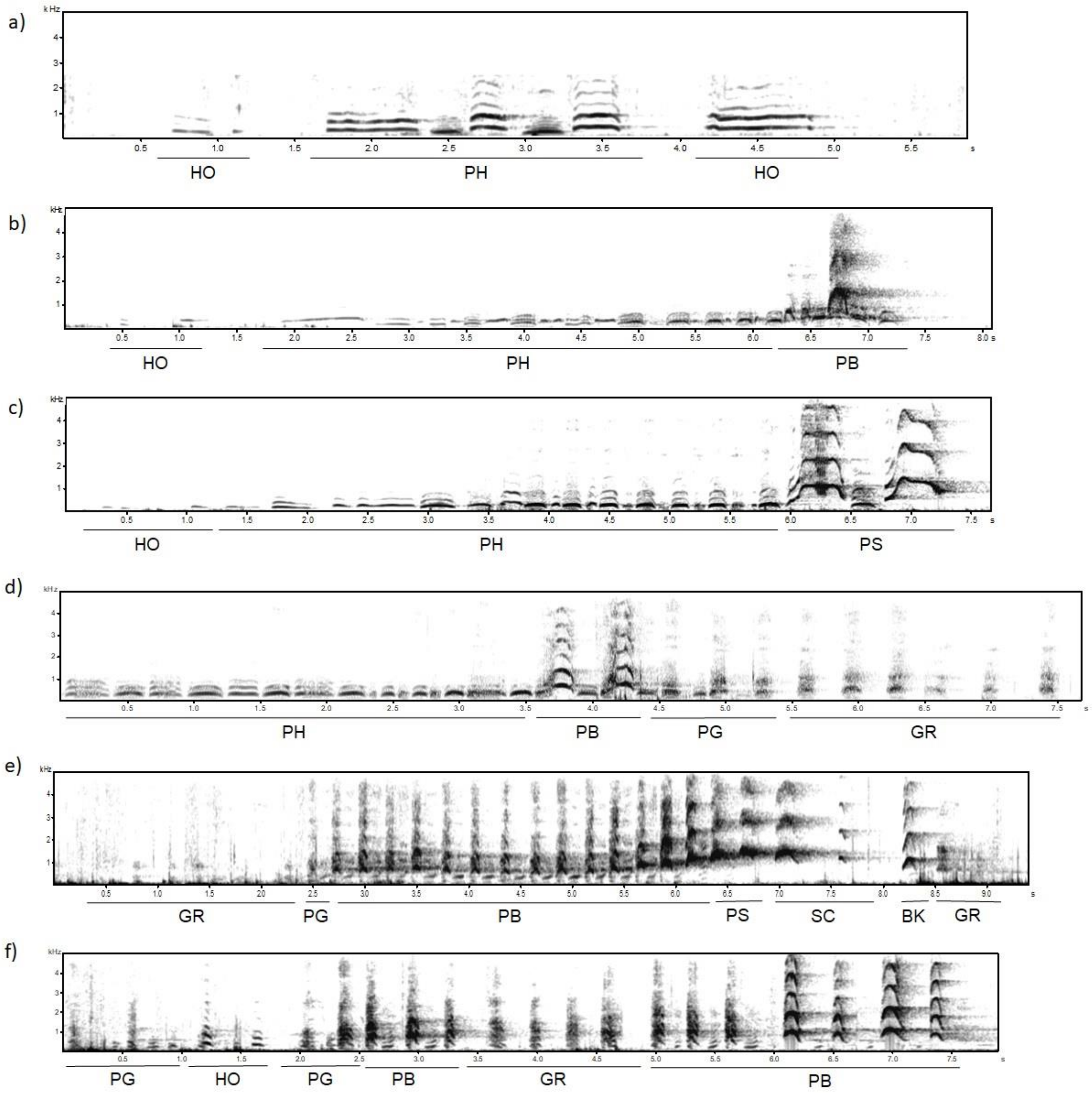


**Figure S3: An example showing the construction of chimpanzee sequences.** Frequency (kHz) in y-axis and time (seconds) in x-axis. Spectrograms (a), (b) and (c) show 3-unit sequences composed by Hoos (single hoo in (a) or series or hoos in (b) and (c)) and series of Panted hoos followed by either a Hoo (a), series of Panted barks (b) or series of Panted screams (c). Spectrogram (d) shows a four-unit sequence composed by series of: Panted hoos, Panted barks, Panted grunts and finally Grunts. Spectrograms (e) and (f) show different long sequences without the structure of the classic Pant-hoot call. The sequence in (e) is composed of Grunts, a single Panted Grunt, Panted barks, Panted screams, Screams, a single

Bark and Grunts. Sequence (f) is composed of Panted grunts, Hoos, Panted grunts, Panted barks, Grunts, and Panted barks.

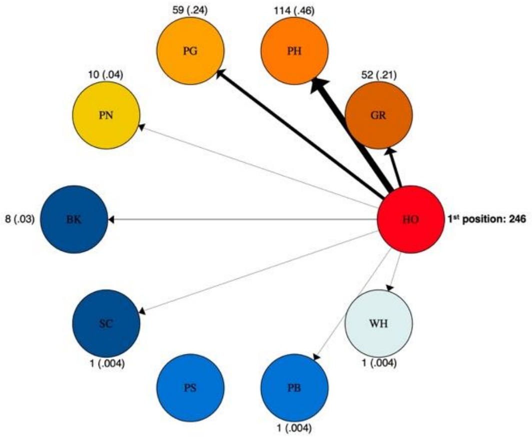


**Figure S4.** Example of two-unit sequences given a certain call in position one (HO). Colours represent the number of times a certain unit is found in the two-unit utterance sample (red-to-turquoise). The size of the directional edges (arrows) expresses the number of times that specific two-unit utterance is found in the study (thick-to-thin). The value on each calls expresses the number of times that the call is found in position two with HO in position one, with corresponding percent value.

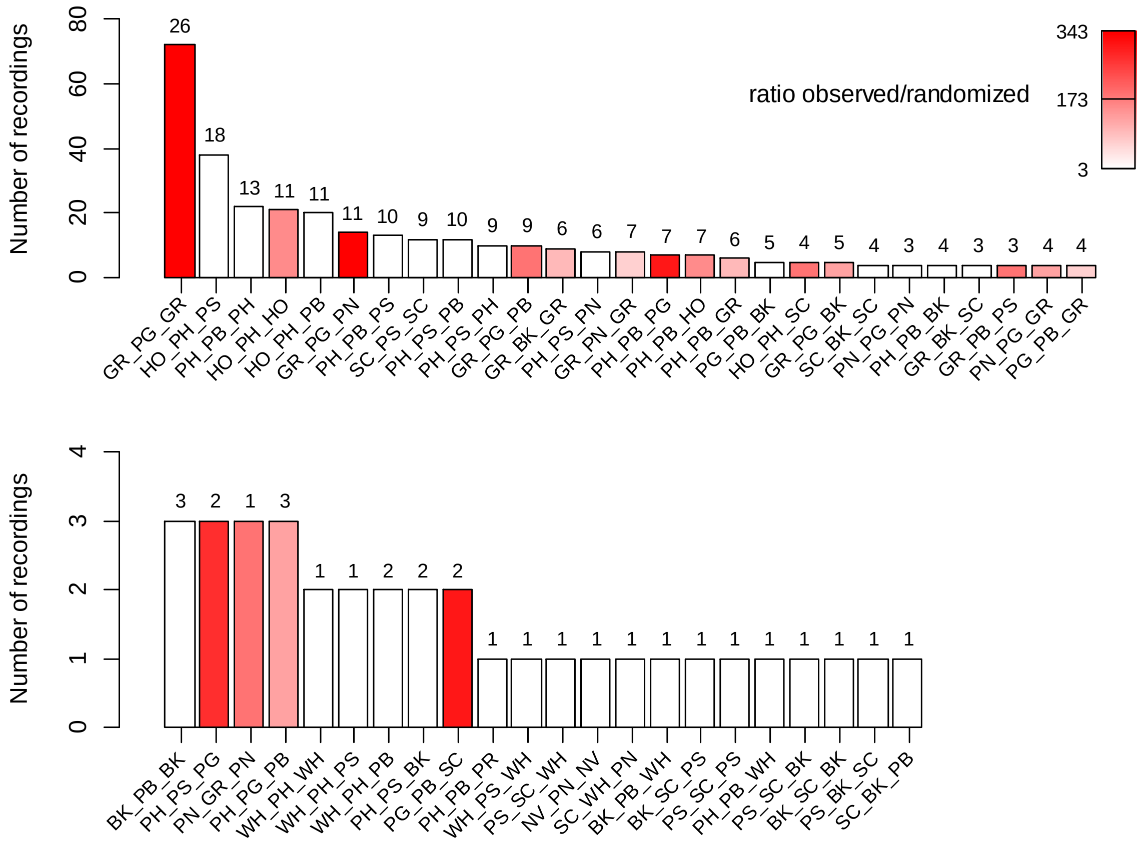


**Figure S5**: Frequency of production of sequences with three vocal units that are produced above chance (i.e., 95% more likely than by random juxtaposition of single vocal units). The height of each bar corresponds to the number of time each sequence was recorded. The colour gradient in the bars depicts the number of time each sequence was observed divided by the number of times each sequence was present on average in each randomization (averaged over 1000 randomizations). The colour range from the lowest in white (i.e., the sequence was present in the randomization only three times less than the number of times it was observed) to the highest ratio in red (i.e., the sequence in randomization 343 times less than the frequency at which it was observed). The number on top of each bar indicates the number of individuals that produced each sequence. The abbreviation for the call names (or single vocal units) are as in Table 1.

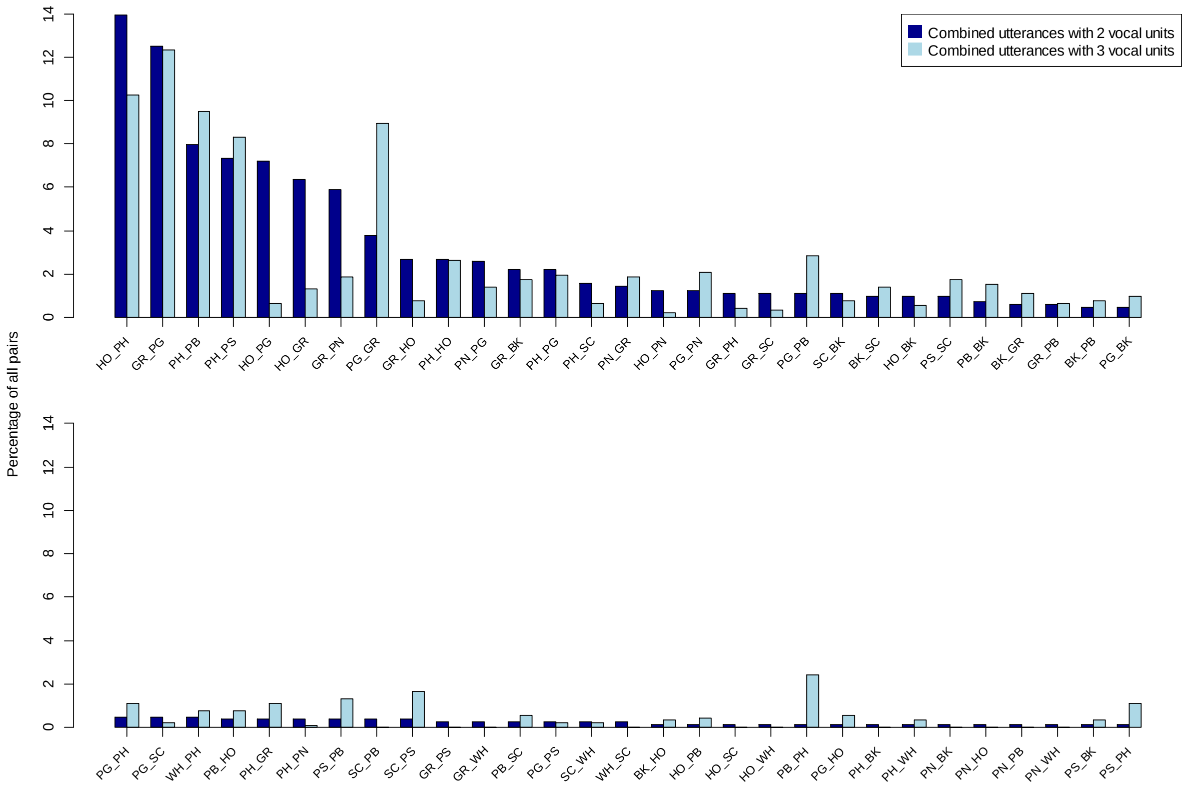


**Figure S6**: Frequency distribution of pairs in sequences with two vocal units (dark blue) and of pairs within sequences with three vocal units (light blue). The height of the bars indicates the percentage of occurrence of each pair within each frequency distribution as a percentage of all pairs recorded. The abbreviations for the call names are as in Table 1.

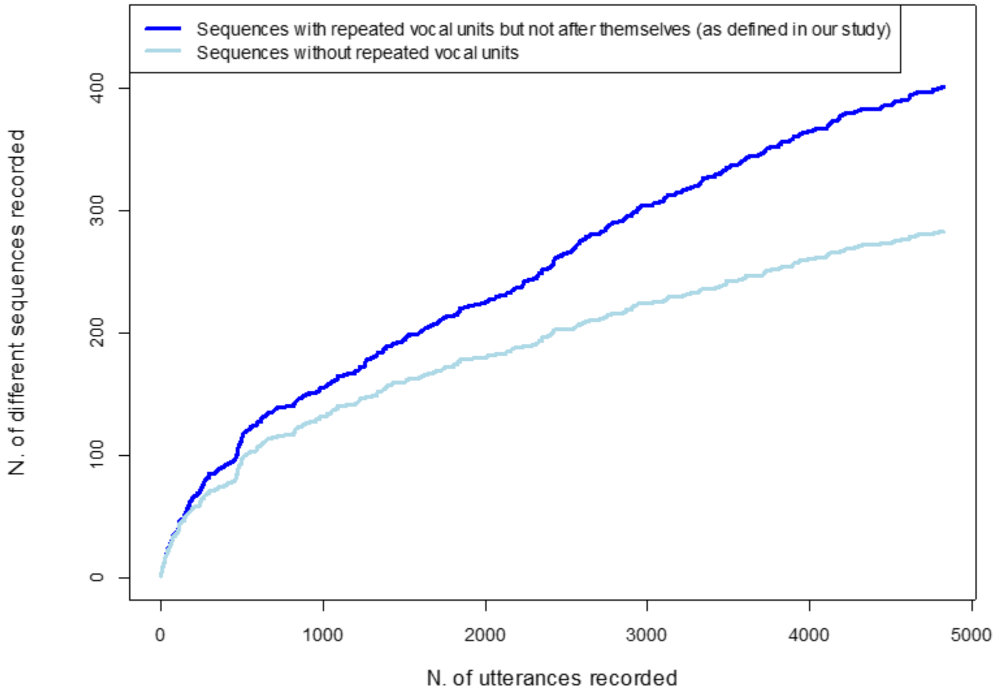


**Figure S7**: Number of different vocal sequences found in the Taï chimpanzee vocal repertoire as a function of the number of utterances recorded.

The dark blue line depicts the number of unique sequences as defined in our study. The sequence considered are the ones where the same vocal unit can occur more than ones within the sequence but not after itself. For instance, A_A_A_B_C would be coded as A_B_C and be no different to a A_B_C sequence. In contrast, A_B_C_A would be different from A_B_C or from B_C_A since A appears twice in the sequence but not after itself.

The light blue line depicts the number of unique sequences in which the same vocal unit is not repeated at all. For instance, the sequence A_B_A_C would be considered the same as the sequence A_B_C since the repetition of A in the first sequence would not be taken into account).

We depict here these two quantification methods since both methods are used in the primate vocal communication literature to assess the diversity of sequences produced by other primate species.

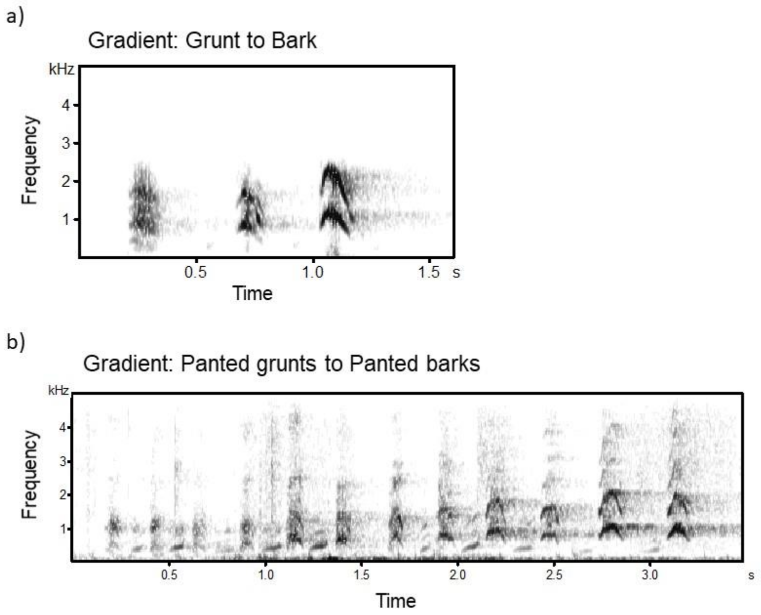


**Figure S8:** Spectrograms of chimpanzees´ graded vocal system showing (a) a gradient from a Grunt to a Bark; and (b) a gradient from Panted-grunts to Panted-barks.

### Supplementary Tables

**Table S1: Details of single utterances and the sequences of different length recorded.**

| Number of vocal units in the utterance | Number of recordings | Number of unique utterance type | List of the unique utterance |
|---|---|---|---|
| 1 | 3242 | 11 | BK, GR, HO, NV, PB, PG, PH, PN, PS, SC, WH |
| 2 | 817 | 58 | BK_GR, BK_HO, BK_PB, BK_SC, GR_BK, GR_HO, GR_PB, GR_PG, GR_PH, GR_PN, GR_PS, GR_SC, GR_WH, HO_BK, HO_GR, HO_PB, HO_PG, HO_PH, HO_PN, HO_SC, HO_WH, PB_BK, PB_HO, PB_PH, PB_SC, PG_BK, PG_GR, PG_HO, PG_PB, PG_PH, PG_PN, PG_PS, PG_SC, PH_BK, PH_GR, PH_HO, PH_PB, PH_PG, PH_PN, PH_PS, PH_SC, PH_WH, PN_BK, PN_GR, PN_HO, PN_PB, PN_PG, PN_WH, PS_BK, PS_PB, PS_PH, PS_SC, SC_BK, SC_PB, SC_PS, SC_WH, WH_PH, WH_SC |
| 3 | 458 | 104 | BK_PB_BK, BK_PB_WH, BK_PH_PB, BK_SC_BK, BK_SC_PS, GR_BK_GR, GR_BK_PB, GR_BK_SC, GR_HO_GR, GR_PB_PS, GR_PB_SC, GR_PG_BK, GR_PG_GR, GR_PG_PB, GR_PG_PH, GR_PG_PN, GR_PG_PS, GR_PG_SC, GR_PH_GR, GR_PH_HO, GR_PH_PB, GR_PN_GR, GR_PN_NV, GR_PN_PG, GR_SC_GR, HO_BK_HO, HO_BK_PS, HO_BK_SC, HO_GR_HO, HO_GR_PG, HO_GR_PN, HO_PB_BK, HO_PB_PG, HO_PB_PS, HO_PG_GR, HO_PG_HO, HO_PG_PH, HO_PG_PN, HO_PH_GR, HO_PH_HO, HO_PH_PB, HO_PH_PG, HO_PH_PS, HO_PH_SC, HO_PH_WH, HO_PN_GR, HO_PN_PG, NV_PN_NV, PG_BK_GR, PG_BK_SC, PG_GR_PG, PG_PB_BK, PG_PB_GR, PG_PB_PG, PG_PB_SC, PG_PH_GR, PG_PH_PB, PH_GR_PG, PH_GR_PN, PH_PB_BK, PH_PB_GR, PH_PB_HO, PH_PB_PG, PH_PB_PH, PH_PB_PR, PH_PB_PS, PH_PB_SC, PH_PB_WH, PH_PG_BK, PH_PG_GR, PH_PG_HO, PH_PG_PB, PH_PG_PH, PH_PG_PS, PH_PG_SC, PH_PS_BK, PH_PS_PB, PH_PS_PG, PH_PS_PH, PH_PS_PN, PH_PS_SC, PH_SC_GR, PN_GR_BK, PN_GR_PG, PN_GR_PN, PN_GR_SC, PN_PG_GR, PN_PG_PH, PN_PG_PN, PN_PH_PN, PS_BK_SC, PS_SC_BK, PS_SC_PS, PS_SC_WH, SC_BK_PB, SC_BK_SC, SC_PS_GR, SC_PS_SC, SC_WH_PN, WH_PH_HO, WH_PH_PB, WH_PH_PS, WH_PH_WH, WH_PS_WH |
| 4 | 170 | 106 | BK_PN_PB_PH, BK_PS_BK_PN, BK_PS_PB_PG, BK_SC_BK_GR, BK_SC_BK_PB, BK_SC_PS_BK, GR_BK_SC_PS, GR_HO_PH_GR, GR_PB_BK_GR, GR_PB_HO_BK, GR_PB_PS_BK, GR_PG_BK_GR, GR_PG_BK_SC, GR_PG_GR_BK, GR_PG_GR_HO, GR_PG_GR_PG, GR_PG_GR_SC, GR_PG_PB_BK, GR_PG_PB_GR, GR_PG_PB_PG, GR_PG_PB_SC, GR_PG_PH_GR, GR_PG_PH_PB, GR_PG_PH_PG, GR_PG_PN_GR, GR_PG_PS_SC, GR_PG_SC_BK, GR_PH_HO_GR, GR_PH_PG_GR, GR_PN_GR_PG, GR_PN_PG_GR, GR_SC_BK_GR, HO_PB_BK_GR, HO_PB_PS_SC, HO_PH_HO_GR, HO_PH_HO_PG, HO_PH_HO_PN, HO_PH_PB_BK, HO_PH_PB_GR, HO_PH_PB_HO, HO_PH_PB_PG, HO_PH_PB_PH, HO_PH_PB_PS, HO_PH_PG_GR, HO_PH_PG_PB, HO_PH_PG_PN, HO_PH_PS_GR, HO_PH_PS_PB, HO_PH_PS_PG, HO_PH_PS_PH, HO_PH_PS_PN, HO_PH_PS_SC, HO_PH_WH_SC, HO_PS_BK_GR, HO_PS_PB_PS, PB_GR_PG_BK, PG_GR_PG_PB, PG_GR_PG_PN, PG_GR_PN_SC, PG_PB_BK_GR, PG_PB_PG_GR, PG_PB_PS_GR, PG_PB_PS_SC, PG_PH_GR_HO, PG_PH_PB_BK, PG_SC_BK_SC, PH_HO_PH_HO, PH_PB_HO_PG, PH_PB_PG_HO, PH_PB_PG_PN, PH_PB_PH_GR, PH_PB_PH_PG, PH_PB_PH_PS, PH_PB_PN_PS, PH_PB_PS_HO, PH_PB_PS_PB, PH_PB_PS_PG, PH_PB_PS_PH, PH_PG_BK_PG, PH_PG_PB_BK, PH_PG_PH_GR, PH_PS_PH_PB, PH_PS_PH_PS, PN_GR_PG_PN, PN_GR_PG_WH, PN_PG_PB_PN, PN_PG_PN_GR, PN_PG_PN_HO, PN_PG_PN_PG, PN_PH_PG_PN, PS_PB_HO_PH, SC_BK_PB_BK, SC_BK_PB_GR, SC_BK_PS_SC, SC_BK_SC_BK, SC_GR_SC_GR, SC_PN_SC_PN, SC_PS_BK_SC, WH_GR_SC_PS, WH_HO_PH_PB, WH_PH_PB_HO, WH_PH_PS_PB, WH_PH_PS_SC, WH_PS_BK_SC, WH_SC_PN_WH, WH_SC_PS_SC |

| | | | |
|---|---|---|---|
| 5 | 90 | 73 | BK_SC_PS_PB_BK, GR_BK_PB_BK_PB, GR_HO_GR_PH_GR, GR_HO_PB_PG_PN, GR_HO_PH_PB_GR, GR_HO_PH_PB_PG, GR_HO_PH_PB_PH, GR_PB_BK_PB_PG, GR_PG_GR_PB_GR, GR_PG_GR_PG_GR, GR_PG_GR_PG_SC, GR_PG_PB_BK_GR, GR_PG_PB_BK_SC, GR_PG_PB_GR_SC, GR_PG_PB_PG_GR, GR_PG_PB_PG_PN, GR_PG_PB_PG_SC, GR_PG_PB_PS_BK, GR_PG_PB_PS_SC, GR_PG_PB_SC_BK, GR_PG_PH_PB_PH, GR_PG_PH_PG_PB, GR_PH_GR_SC_GR, GR_PH_PB_BK_GR, GR_PH_PB_PH_GR, GR_PH_PB_PS_SC, GR_PH_PG_PN_GR, GR_PN_BK_PB_BK, GR_PN_GR_PG_GR, GR_PN_GR_PG_PN, GR_PN_PG_PB_HO, GR_SC_PS_SC_BK, HO_GR_PG_PB_PG, HO_PB_PS_GR_BK, HO_PG_PB_PG_PB, HO_PH_GR_HO_PB, HO_PH_HO_PH_HO, HO_PH_PB_GR_PG, HO_PH_PB_PG_GR, HO_PH_PB_PG_PB, HO_PH_PB_PH_GR, HO_PH_PB_PH_HO, HO_PH_PB_PS_SC, HO_PH_PN_PG_PB, HO_PH_PS_PB_PS, PG_GR_PG_PB_PG, PG_PB_BK_SC_PB, PG_PB_BK_SC_PS, PG_PB_PG_PB_GR, PG_PB_PG_PB_PG, PG_PB_SC_BK_GR, PG_PH_PB_BK_PH, PH_GR_PG_PB_BK, PH_PB_PH_PB_PH, PH_PB_PH_PS_PG, PH_PB_PS_PB_PG, PH_PB_PS_PH_HO, PH_PB_SC_BK_SC, PH_PG_PB_PS_SC, PH_PG_PH_PB_PG, PH_PS_PB_PS_BK, PN_GR_PG_GR_PN, PN_GR_PG_PB_PG, PN_GR_PN_GR_PG, PN_GR_PN_PG_GR, PN_PG_PB_PG_PH, PN_PG_PB_PG_PN, PS_PB_BK_PS_BK, PS_SC_PS_SC_BK, SC_BK_PB_SC_BK, SC_BK_SC_BK_PS, SC_PS_SC_PS_SC, WH_PH_WH_PH_WH |
| 6 | 29 | 29 | BK_PS_BK_SC_BK_SC, GR_BK_GR_PG_BK_GR, GR_BK_PS_BK_SC_PB, GR_BK_SC_GR_BK_SC, GR_HO_GR_HO_PH_PS, GR_HO_PH_PG_PB_BK, GR_PG_GR_HO_PH_GR, GR_PG_GR_HO_PH_HO, GR_PG_GR_PG_GR_PG, GR_PG_GR_PG_HO_GR, GR_PG_GR_SC_PS_SC, GR_PG_HO_PG_PB_GR, GR_PG_PB_GR_PB_PG, GR_PG_PB_PG_PB_PN, GR_PG_PB_PS_SC_BK, GR_PG_PB_SC_BK_GR, GR_PH_PB_PS_PB_PS, GR_SC_PS_SC_BK_SC, HO_GR_PG_GR_PH_HO, HO_PH_PB_PG_GR_PN, HO_PH_PB_PH_PB_GR, HO_PH_PS_PB_PS_PH, PG_GR_PG_PB_BK_GR, PH_PB_PG_PH_PG_GR, PH_PB_PG_PS_PG_GR, PH_PG_PB_GR_PG_PB, PN_GR_PN_GR_PN_PG, PN_PG_PB_PG_PB_BK, PN_PG_PH_PB_PG_GR" |
| 7 | 12 | 12 | GR_BK_GR_BK_GR_BK_GR, GR_PG_GR_PG_GR_PG_PN, GR_PG_PB_PS_PB_PS_SC, GR_PG_PH_HO_GR_PG_GR, GR_PG_SC_PS_SC_BK_SC, HO_PH_PB_PH_PB_PB_BK, HO_PH_PB_PH_PB_PH_HO, HO_PH_PB_PH_PS_PH_PB, HO_PH_PG_PH_PG_GR_PG, HO_PN_GR_PH_PG_GR_PG, PH_PB_PH_PB_PH_PB_PG, PH_PB_PH_PB_PH_PB_PH" |
| 8 | 4 | 4 | GR_BK_WH_SC_PS_GR_WH_SC, GR_PG_PB_PG_GR_PG_PB_GR, GR_PG_PB_PG_PB_PG_PB_BK, HO_PH_GR_PG_PB_PG_PB_PG |
| 9 | 2 | 2 | BK_GR_PG_PB_GR_GR_BK_GR_BK, HO_PH_PS_PH_PS_HO_PH_PS_SC |
| 10 | 2 | 2 | GR_BK_PS_SC_BK_SC_BK_PB_BK_PB, PG_GR_PN_GR_PG_PN_PG_PB_GR_PN |
| **Total** | **4826** | **401** | |

**Table S2: Positional occurrences in two-unit sequences**

second position (columns); first position (rows)

| unit | BK | GR | HO | NV | PB | PG | PH | PN | PR | PS | SC | WH | total |
|---|---|---|---|---|---|---|---|---|---|---|---|---|---|
| BK | | 5 | 1 | 0 | 4 | 0 | 0 | 0 | 0 | 0 | 8 | 0 | 18 |
| GR | 18 | | 22 | 0 | 5 | 102 | 9 | 48 | 0 | 2 | 9 | 2 | 217 |
| HO | 8 | 52 | | 0 | 1 | 59 | 114 | 10 | 0 | 0 | 1 | 1 | 246 |
| NV | 0 | 0 | 0 | | 0 | 0 | 0 | 0 | 0 | 0 | 0 | 0 | 0 |
| PB | 6 | 0 | 3 | 0 | | 0 | 1 | 0 | 0 | 0 | 2 | 0 | 12 |
| PG | 4 | 31 | 1 | 0 | 9 | | 4 | 10 | 0 | 2 | 4 | 0 | 65 |
| PH | 1 | 3 | 22 | 0 | 65 | 18 | | 3 | 0 | 60 | 13 | 1 | 186 |
| PN | 1 | 12 | 1 | 0 | 1 | 21 | 0 | | 0 | 0 | 0 | 1 | 37 |
| PR | 0 | 0 | 0 | 0 | 0 | 0 | 0 | 0 | | 0 | 0 | 0 | 0 |
| PS | 1 | 0 | 0 | 0 | 3 | 0 | 1 | 0 | 0 | | 8 | 0 | 13 |
| SC | 9 | 0 | 0 | 0 | 3 | 0 | 0 | 0 | 0 | 3 | | 2 | 17 |
| WH | 0 | 0 | 0 | 0 | 0 | 0 | 4 | 0 | 0 | 0 | 2 | | 6 |
| total | 48 | 103 | 50 | 0 | 91 | 200 | 133 | 71 | 0 | 67 | 47 | 7 | 817 |

**Table S3: Positional occurrences in three-unit sequences**

| two-unit | Head | Tail | total |
|---|---|---|---|
| GR_PG | 105 | 8 | 113 |
| HO_PH | 94 | 0 | 94 |
| PH_PB | 62 | 25 | 87 |
| PH_PS | 36 | 40 | 76 |

**Table S4: Transitional relationships (forward) in three-unit sequences**

Two-unit utterance (columns); following call (rows)

| unit | GR_PG | HO_PH | PH_PB | PH_PS |
|---|---|---|---|---|
| BK | 5 | 0 | 4 | 2 |
| GR | 72 | 6 | 6 | 0 |
| HO | 0 | 21 | 7 | 0 |
| NV | 0 | 0 | 0 | 0 |
| PB | 10 | 20 | 0 | 12 |
| PG | 0 | 3 | 7 | 3 |
| PH | 2 | 0 | 22 | 10 |
| PN | 14 | 0 | 0 | 8 |
| PR | 0 | 0 | 1 | 0 |
| PS | 1 | 38 | 13 | 0 |
| SC | 1 | 5 | 1 | 1 |
| WH | 0 | 1 | 1 | 0 |
| total | 105 | 94 | 62 | 36 |

**Table S5: Transitional relationships (backward) in three-unit sequences**

Two-unit utterance

| preceding call | unit | GR_PG | HO_PH | PH_PB | PH_PS |
|---|---|---|---|---|---|
| | BK | 0 | 0 | 1 | 0 |
| | GR | 0 | 0 | 1 | 0 |
| | HO | 3 | 0 | 20 | 38 |
| | NV | 0 | 0 | 0 | 0 |
| | PB | 0 | 0 | 0 | 0 |
| | PG | 2 | 0 | 1 | 0 |
| | PH | 1 | 0 | 0 | 0 |
| | PN | 2 | 0 | 0 | 0 |
| | PR | 0 | 0 | 0 | 0 |
| | PS | 0 | 0 | 0 | 0 |
| | SC | 0 | 0 | 0 | 0 |
| | WH | 0 | 0 | 2 | 2 |
| | total | 8 | 0 | 25 | 40 |
